# CDK1 couples proliferation with protein synthesis

**DOI:** 10.1101/816850

**Authors:** Katharina Haneke, Johanna Schott, Doris Lindner, Anne K. Hollensen, Christian K. Damgaard, Cyril Mongis, Michael Knop, Wilhelm Palm, Alessia Ruggieri, Georg Stoecklin

## Abstract

Cell proliferation exerts a high demand on protein synthesis, yet the mechanisms coupling the two processes are not fully understood. A kinase and phosphatase screen for activators of translation, based on the formation of stress granules in human cells, revealed cell cycle-associated kinases as major candidates. CDK1 was identified as a positive regulator of global translation, and cell synchronization experiments showed that this is an extra-mitotic function of CDK1. Dephosphorylation of eIF2α and S6K1 signaling were found to act downstream of CDK1. Moreover, Ribo-Seq analysis uncovered that CDK1 exerts a particularly strong effect on the translation of 5’TOP mRNAs, which includes mRNAs encoding for ribosomal proteins and several translation factors. This effect requires the 5’TOP mRNA-binding protein LARP1, concurrent to our finding that LARP1 phosphorylation is strongly dependent on CDK1. Taken together, our results show that CDK1 provides a direct means to couple cell proliferation with biosynthesis of the translation machinery and the rate of protein synthesis.

## INTRODUCTION

Cell growth, proliferation and progression through the cell cycle strongly depend on the synthesis of new proteins (Pardee, 1989; Polymenis and Aramayo, 2015). On the one hand, cells exert temporal control over the production of specific proteins during the different phases of the cell cycle (Aviner et al., 2013; Stumpf et al., 2013; Tanenbaum et al., 2015). On the other hand, cells also need to adjust the overall rate of protein synthesis to the proliferation rate in order to maintain cell size and functionality (Foster et al., 2010). It is therefore not surprising that modifications of the translation machinery can affect cell proliferation rates, and that deregulation of protein synthesis is increasingly recognized as a major driver of cell transformation (Ruggero and Pandolfi, 2003; Silvera et al., 2010; Truitt and Ruggero, 2016)

A few signaling pathways are known to regulate protein synthesis in response to proliferative cues. The mechanistic target of rapamycin complex 1 (mTORC1), e.g., functions as a signaling node that adjusts protein synthesis to cell growth rates and the metabolic status of the cell (Laplante and Sabatini, 2012). mTORC1 directly phosphorylates 4E-BPs, thereby promoting the translation of a distinct group of mRNAs that strongly depend on the eukaryotic translation initiation factor (eIF) 4E (Gandin et al., 2016; Nandagopal and Roux, 2015). mTORC1 further enhances the translation of mRNAs containing a 5’ terminal oligo pyrimidine tract (5’TOP) motif, which includes many mRNAs encoding ribosomal proteins and translation factors (Meyuhas and Kahan, 2015).

The protooncogenes Ras and Myc also control protein synthesis in order to coordinate cellular growth rates with extracellular growth stimuli. While Myc mostly controls translation through transcriptional upregulation of ribosomal components and translation factors (van Riggelen et al., 2010), the Ras/Erk signaling pathway shares some common downstream signals with mTORC1 including phosphorylation of ribosomal protein S6 (RPS6) (Roux and Topisirovic, 2018).

While numerous translation factors are known to be phosphorylated (Roux and Topisirovic, 2018), the regulatory impact of phosphorylation is established only for a few factors such as eIF2α, 4E-BPs and eEF2 (Jackson et al., 2010; Kenney et al., 2014). Ribosomal proteins are also known to carry various posttranslational modifications (Shi and Barna, 2015), yet the role of these modifications in controlling protein synthesis is poorly understood. Recently, a systematic approach to identify translationally relevant phosphorylation sites on ribosomal proteins revealed that phosphorylation of RPL12 controls the translation of mitosis-specific proteins (Imami et al., 2018).

At the core of the cell cycle, cyclin-dependent kinases (CDKs) drive cells through the different phases of the cell cycle. In G1, Cyclin D-CDK4/6 (early) and Cyclin E-CDK2 (late) prepare entry into S-phase, where Cyclin A-CDK2 takes over and orchestrates replication, followed by activation of Cyclin A/B-CDK1 promoting passage through G2 and entry into M-phase (Malumbres and Barbacid, 2005). Interestingly, CDK1 can substitute for the other CDKs and was found to be sufficient for driving the mammalian cell cycle (Santamaria et al., 2007). CDK1 has also been linked to the control of protein synthesis during M-phase (Shuda et al., 2015; Sivan et al., 2011)

In this study, we made use of the fact that a global decrease in translation initiation is coupled to the assembly of cytoplasmic stress granules (SGs), aggregates that arise through phase separation of stalled mRNAs and associated factors from the surrounding cytosol (Kedersha et al., 2013). To identify novel regulators of protein synthesis, we conducted an siRNA screen against all human kinases an phosphatases using SG formation as a visual read-out. Since cell cycle-associated kinases were among the primary candidates identified by the screen, we chose to pursue CDK1 and characterize its role in protein synthesis. Our results demonstrate that CDK1 acts outside of mitosis as a general activator of translation that allows direct adaptation of protein synthesis to the rate of cell proliferation.

## RESULTS

### Identification of kinases and phosphatases suppressing SG assembly

With the aim to identify kinases and phosphatases that enhance global protein synthesis under regular growth conditions, we knocked down 711 human kinases and 256 phosphatases in HeLa cells stably expressing the SG marker GFP-G3BP1, using 4 independent siRNAs for each phosphotransferase. After 72 hours, cells were monitored for the presence of SGs. As expected, GFP-G3BP1 was evenly distributed in the cytoplasm in control knock down (kd) cells (Fig. 1A). SG formation was detected in a small fraction of the kd cultures, and typically occurred only in a subpopulation of cells (examples in Fig. 1A). For every phosphotransferase we calculated a SG score, which reflects both the strength of the phenotype and its reproducibility, and found that kd of 54 kinases (8%) and 15 phosphatases (6%) led to SG formation with a SG score >10 (with at least 2 different siRNAs) or >40 (with 1 siRNA). In comparison, control cells transfected with non-targeting siRNAs had an average SG score of 1.9 (Fig. 1B and 1C, Table S1).

**Figure 1.**
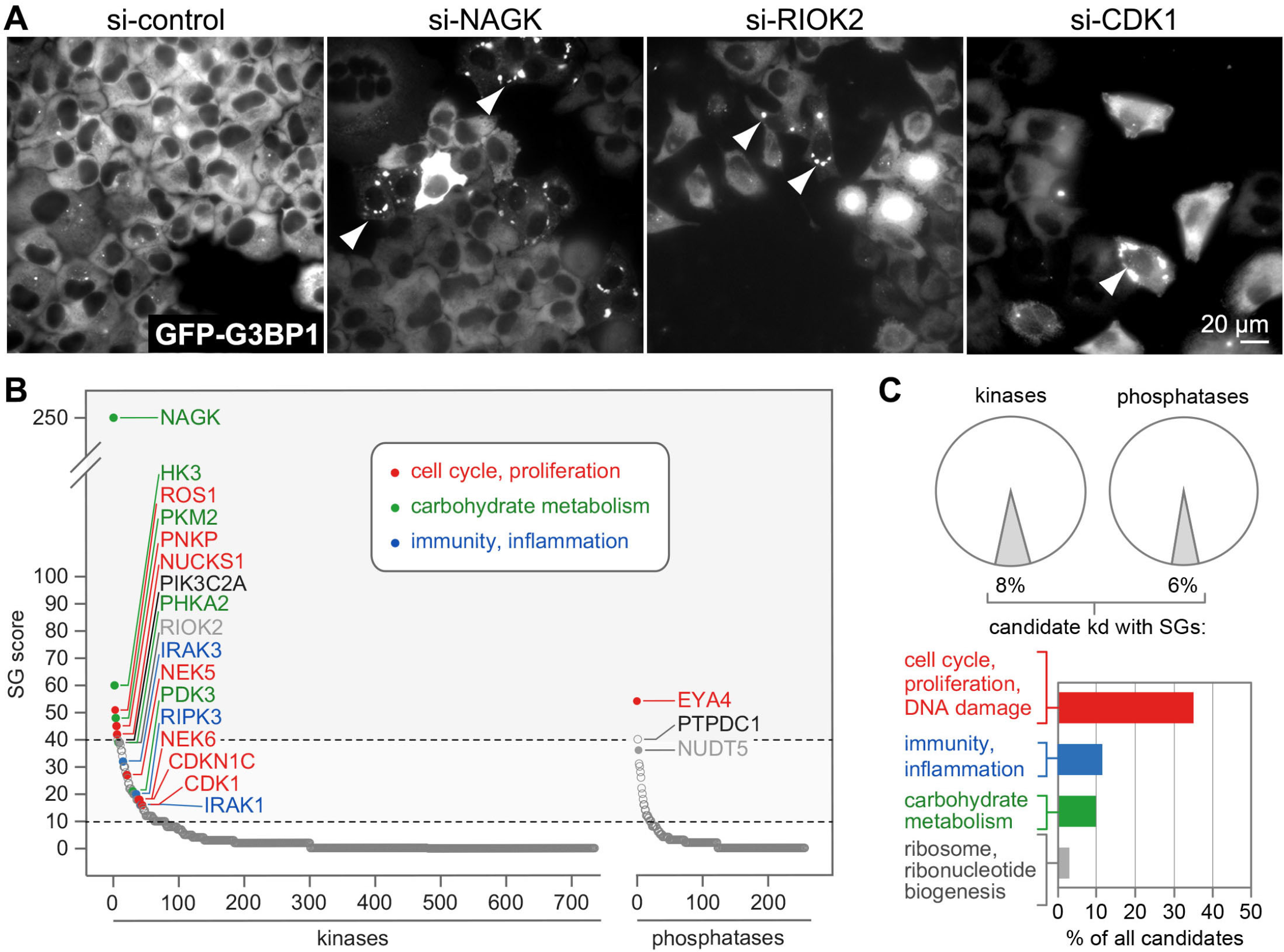
SG assembly screen under regular growth conditions. **(A)** The assembly of SGs was monitored in HeLa cells stably expressing GFP-G3BP1 following kd of 711 kinases and 256 phosphatases. Cells were transfected with 4 individual siRNAs per gene and 72 hours later fixed for fluorescence microscopy. Representative images of the screen are shown; arrowheads indicate SG-containing cells. **(B)** The screen was analyzed by calculating a SG score for each kinase/phosphatase kd, and the result was depicted by sorting all kds according to their SG score. Candidate kinases/phosphatases were identified by a SG score >10 (with at least 2 different siRNAs) or >40 (with 1 siRNA); some of the candidates were labeled in the graph. **(C)** The graph depicts cellular functions highly represented among the candidate kinases/phosphatases, based on functional annotation in NCBI Gene and Uniprot databases.

To our surprise, phosphotransferases associated with cell cycle regulation, proliferation or DNA damage were highly represented among the candidates (35%, Fig. 1C). Those associated with immunity and inflammation (12%) or carbohydrate metabolism (10%) were also abundant, whereas only few candidates were associated with ribosome and ribonucleotide biogenesis (3%). Given its central role for mitotic entry and its general importance in the cell cycle (Itzhaki et al., 1997; Santamaria et al., 2007), we decided to pursue CDK1 as a candidate that may connect proliferation rates with global protein synthesis.

### Inhibition of CDK1 reduces protein synthesis

Since SG-based screens not only report on regulators of translation, but also on downstream factors that control the assembly and disassembly of SGs, it was important to test if CDK1 influences global translation rates. To this end, we treated HeLa cells for 1 to 24 hours with the selective, ATP-competitive CDK1 inhibitor Ro3306 (Vassilev et al., 2006). CDK1 inhibition (CDKi) led to the assembly of SGs (Fig. 2A), and by polysome profile analysis we observed a progressive decrease in the percentage of polysomal ribosomes (Fig. 2B), a measure that reflects the proportion of ribosomes engaged in translation. We also quantified polypeptide synthesis using a puromycin incorporation assay, and found a similar time-dependent decrease in response to Ro3306 treatment (Fig. 2C and 2D). These results could be confirmed using a less selective CDK inhibitor, Roscovitine (Cicenas et al., 2015) (Fig. S1A).

**Figure 2.**
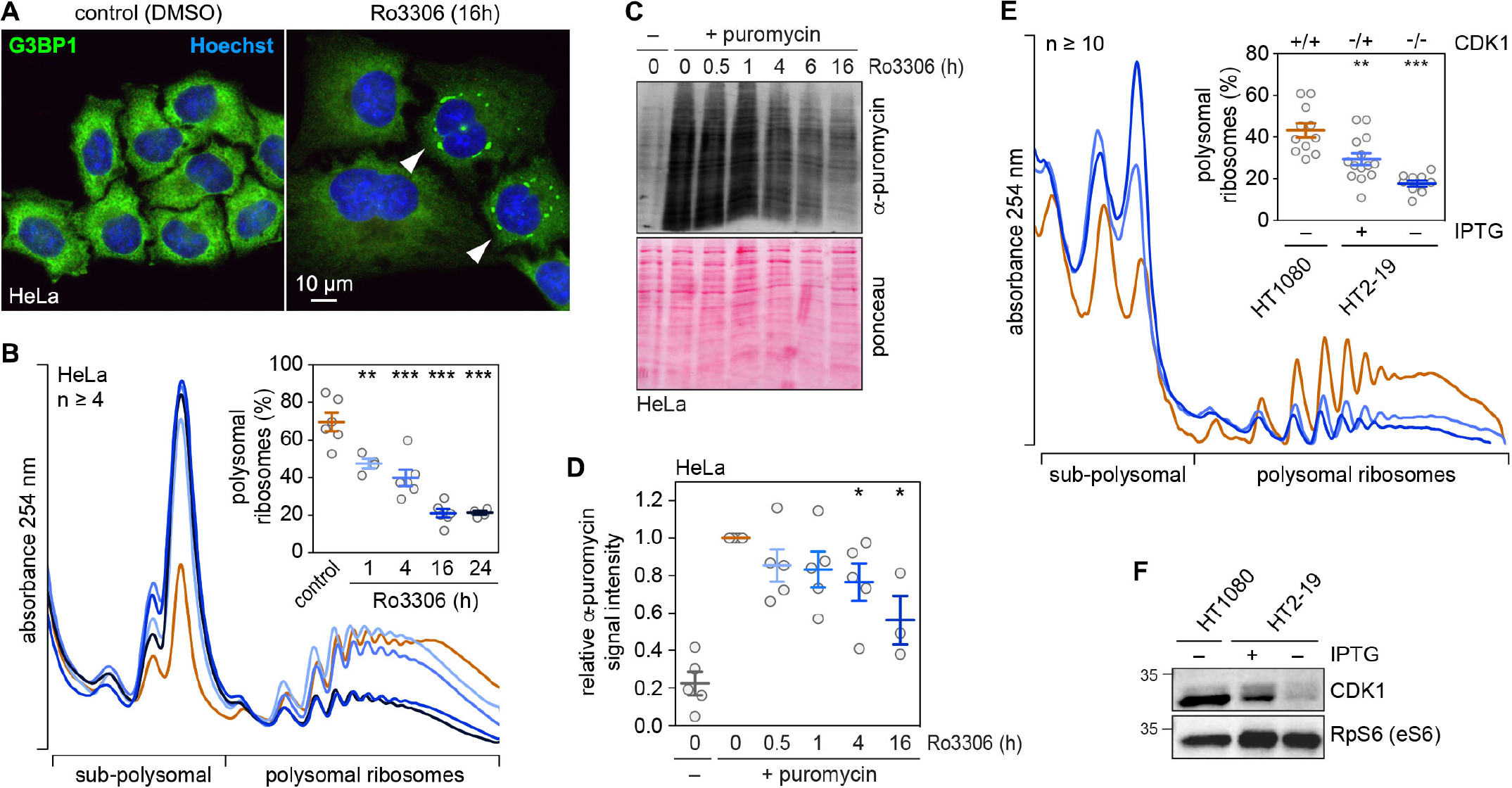
Global translation suppression after pharmacological or genetic CDK1 inhibition. **(A)** HeLa cells were treated either with solvent (DMSO) or the CDK1 inhibitor Ro3306 (10 μM) for 16 h. SG formation was analyzed by IF microscopy of fixed cells stained with anti-G3BP1 antibody and Hoechst. **(B)** Polysome profiles from DMSO- or Ro3306-treated HeLa cells were recorded after sucrose density gradient centrifugation; the percentage of polysomal ribosomes is represented in the inset (average ± SEM, n ≥ 4). Statistical significance was determined by unpaired Student’s t test; **, p ≤ 0.01; ***, p ≤ 0.001. **(C)** Incorporation of puromycin into nascent polypeptides was analyzed by SDS-PAGE and Western blotting. Puromycin-labeled polypeptides were detected with anti-puromycin antibody; ponceau staining served as loading control. **(D)** Puromycin incorporation signal intensities were normalized to the ponceau staining and values were calculated relative to DMSO-treated control samples (average ± SEM, n ≥ 3). Statistical significance was determined by one-sample Student’s t test; *, p ≤ 0.05. **(E)** HT1080 and HT2-19 cells were seeded at sub-confluency and kept in the presence or absence of IPTG (0.2 mM) for 7 days. Polysome profiles were recorded; the percentage of polysomal ribosomes is represented in the inset (average ± SEM, n ≥ 10). Statistical significance was determined by unpaired Student’s t test; **, p ≤ 0.01; ***, p ≤ 0.001. **(F)** CDK1 expression in HT1080 and HT2-19 cells was assessed by Western blot analysis; RPS6 levels served as loading control.

We then sought genetic evidence for a role of CDK1 in controlling protein synthesis. Since CDK1 is an essential gene, we made use of HT2-19, a human HT1080-derived cell line that contains one inactivated CDK1 allele, whereas the other allele is under control of a lac repressor and hence transcribed only in the presence of IPTG (Itzhaki et al., 1997). CDK1 levels were reduced at least 2-fold in HT2-19 cells cultured in presence of IPTG, and polysomal ribosomes decreased from 43% in parental HT1080 to 32% (Fig. 2E and 2F). CDK1 became barely detectable when HT2-19 cells were kept in the absence of IPTG for 7 days, and polysomal ribosomes dropped further to 18% (Fig. 2E and 2F). These cells did not divide anymore but increased in cell size (Fig. S1B).

### CDK1 controls global translation in a cell cycle-independent manner

CDK1 activity changes throughout the cell cycle: it starts to increase during S-phase, reaches its maximum in metaphase, and declines rapidly in anaphase (Bashir and Pagano, 2005). In line with its activity profile, CDK1 was shown to control translation during mitosis both at the level of translation initiation via phosphorylation of raptor (Ramirez-Valle et al., 2010), 4E-BP1 (Heesom et al., 2001; Shuda et al., 2015; Velasquez et al., 2016), S6K1 (Papst et al., 1998; Shah et al., 2003) and eIF4GI (Dobrikov et al., 2014), as well as at the level of elongation via phosphorylation of eEF1B (Monnier et al., 2001; Sivan et al., 2011) and eEF2K (Smith and Proud, 2008).

We noted that CDK1i led to SG formation only in about 10% of cells, which might be related to the peak of CDK1 activity in mitosis. In order to test if SG formation upon CDKi is restricted to a specific phase of the cell cycle, we made use of the FUCCI system and applied Ro3306 to HeLa cells stably expressing either Kusabira-Orange-Cdt1 or mVenus-Geminin (Sakaue-Sawano et al., 2008). While Cdt1 is expressed during G1- and early S-phase, Geminin is expressed in S-phase, G2-phase and mitosis. Quantification of SG-positive cells revealed no preference for a particular cell cycle phase since 10% of the Kusabira-Orange-Cdt1-positive and 9% of the mVenus-Geminin-positive cells contained SGs upon CDKi (Fig. 3A and 3B). This result suggested that CDK1 enhances global protein synthesis in a cell cycle phase-independent manner.

**Figure 3.**
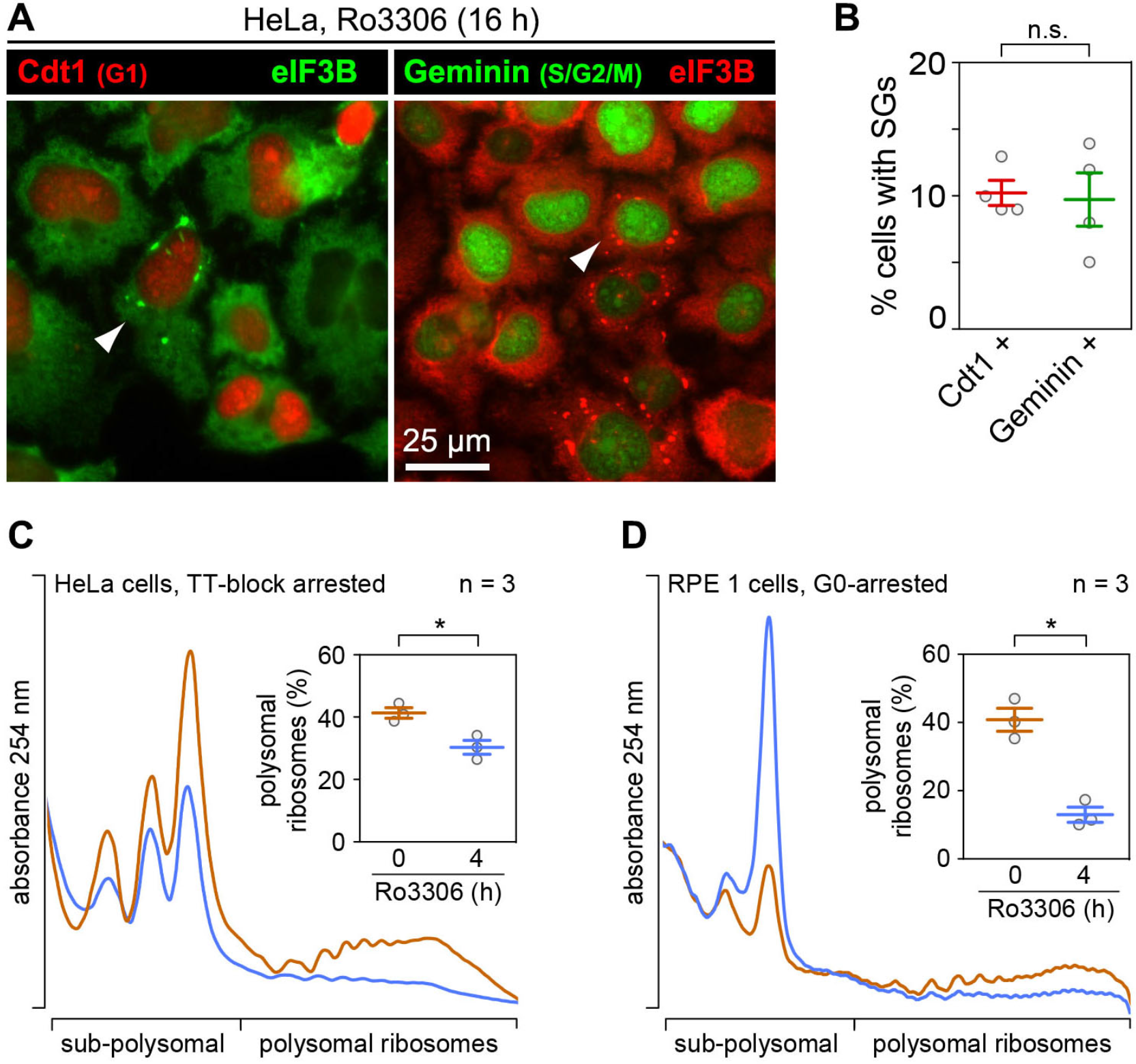
Cell cycle phase-independent translation suppression upon CDK1 inhibition. **(A)** HeLa FUCCI cells were treated with Ro3306 (10 μM) for 16 h, fixed and analyzed for SG formation by IF microscopy upon staining with anti-eIF3B antibody. HeLa cells stably expressing Kusabira-Orange-Cdt1 (marker for G1- and early S-phase) were used in the left panel; HeLa cells stably expressing mVenus-Geminin (marker for S-, G2- and M-phase) were used in the right panel. **(B)** Quantification of the percentage of SG-containing cells in Kusabira-Orange-Cdt1-positive (left panel) or mVenus-Geminin-positive cells (right panel, n = 4). **(C)** HeLa cells were arrested in G1 phase by a double thymidine block (TT) and, without release from the block, treated either with solvent (DMSO) or Ro3306 (10 μM) for 4 h. Polysome profiles were recorded; the percentage of polysomal ribosomes is represented in the inset (average ± SEM, n = 3). **(D)** RPE-1 cells were serum-starved for 48 h and subsequently treated with DMSO or Ro3306 (10 μM) for 4 h. Polysome profiles were recorded; the percentage of polysomal ribosomes is represented in the inset (average ± SEM, n = 3). In (B–D), statistical significance was determined by paired Student’s t test; *, p ≤ 0.05.

To further explore this possibility, we arrested HeLa cells in early S-phase using a double thymidine (TT) block and, without release from the block, subjected them to CDKi. Compared to asynchronously proliferating cells (with 70% polysomal ribosomes, Fig. 2B), the cell cycle arrest alone led to a reduction of global protein synthesis (41% polysomal ribosomes), and treatment with Ro3306 for 4 hours caused a further decrease to 30% polysomal ribosomes (Fig. 3C).

Likewise, we tested non-proliferating RPE1 cells after 48 hours of serum starvation. The cells had entered G0-phase, visible through the appearance of primary cilia (Fig. S2), and still responded to CDK1i by a strong reduction of their translation rate (Fig. 3D). From these experiments we concluded that enhancing protein synthesis is an extra-mitotic function of CDK1, which likely serves as a means to adjust protein synthesis to the overall proliferation rate rather than to a specific phase of the cell cycle.

### eIF2α phosphorylation and S6K1 contribute to translation control by CDK1

We then sought to explore the signaling pathway by which CDK1 controls protein synthesis. Various types of stress cause suppression of translation initiation via phosphorylation of eIF2α at serine (S)51, which prevents recharging of the initiator eIF2-GTP-tRNA_i_^Met^ ternary complex (Jackson et al., 2010). Western blot analysis of cytoplasmic lysates from HeLa cells indicated that CDK1i leads to robust phosphorylation of eIF2α after 16 hours of Ro3306 treatment (Fig. 4A and 4B), whereas the onset of translation suppression was visible already 1 hour after CDK1i (Fig. 2B–D). In line with this notion, translation suppression upon Ro3306 treatment was partially impaired in mouse embryonic fibroblasts (MEFs) containing a bi-allelic phospho-deficient eIF2α-S51A (AA) mutation (Scheuner et al., 2001) as compared to MEFs expressing wild-type eIF2α-S51 (SS) alleles (Fig. 4C, S3A and S3B). Thus, we concluded that eIF2α phosphorylation is alone not responsible for, but contributes to translation inhibition after CDK1i.

**Figure 4.**
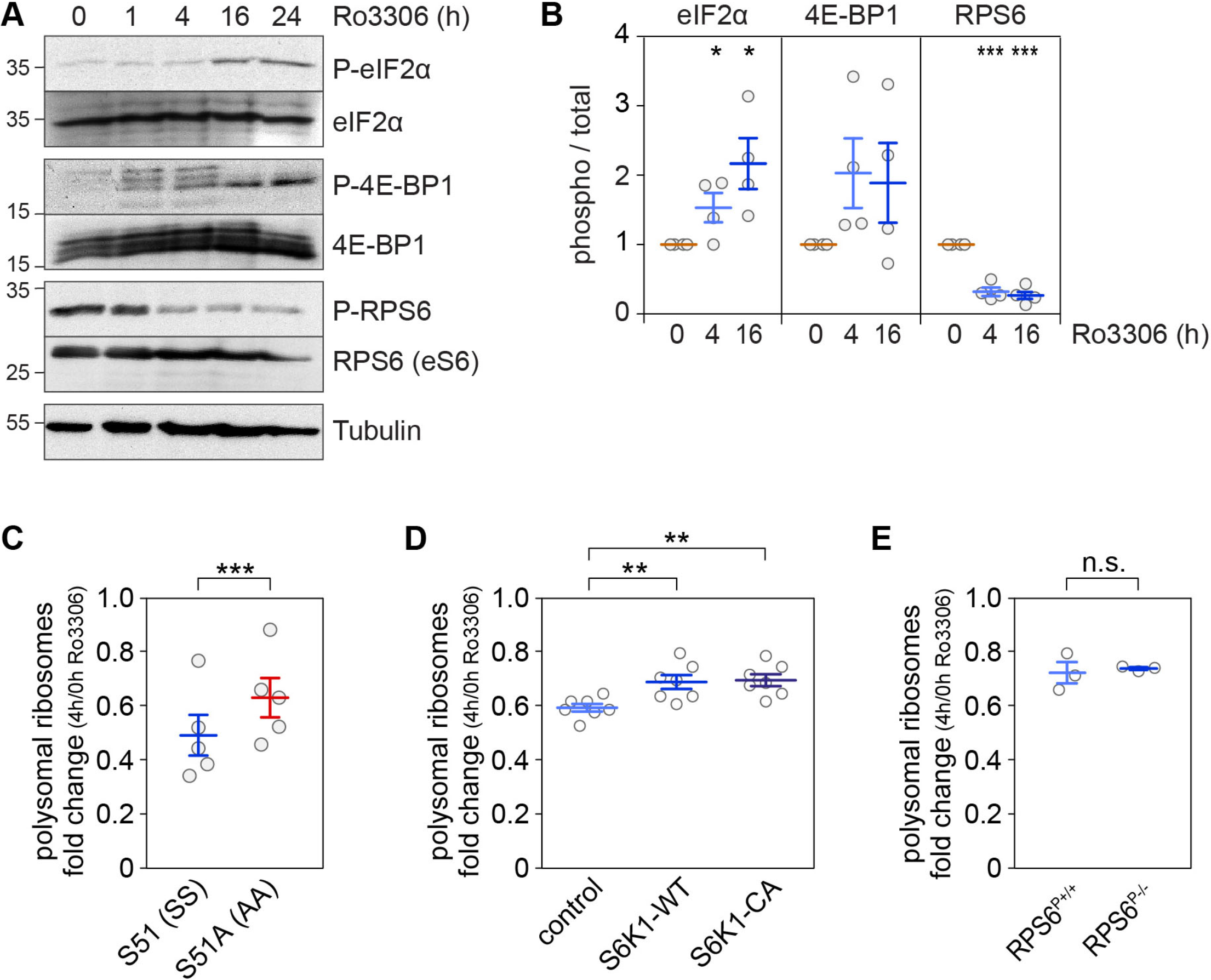
Pathways signaling translational control downstream of CDK1i. **(A)** Protein lysates were prepared from HeLa cells treated with solvent (DMSO, 24 h) or Ro3306 (10 μM, 1–24 h), and the phosphorylation status of eIF2α (S51), RPS6 (eS6) (S235/S236) and 4E-BP1 (T37/T46) were analyzed by Western blot analysis. **(B)** The phosphorylation level of eIF2α (S51), RPS6 (S235/S236) and 4E-BP1 (T37/T46) was quantified from Western blot analyses as shown in (A) (average ± SEM, n = 4). Statistical significance was determined by one-sample Student’s t test; *, p ≤ 0.05; ***, p ≤ 0.001. **(C)** The fold-change in polysomal ribosomes (4 h Ro3306 / DMSO control) was calculated based on polysome profiles recorded from eIF2α WT S51 (SS) and phosphodeficient S51A (AA) MEFs. **(D)** The fold-change in polysomal ribosomes was determined as in (C) from HeLa cells (control), HeLa cells overexpressing HA-S6K1-WT and HeLa cells overexpressing constitutively active HA-S6K1-CA. **(E)** The fold-change in polysomal ribosomes was determined as in (C) from RPS6 WT (RPS6^P+/+^) and RPS6 phosphodeficient S235A, S236A, S240A, S244A, S247A (RPS6^P−/−^) MEFs. In (C–E), statistical significance was determined by paired Student’s t test; **, p ≤ 0.01; ***, p ≤ 0.001.

Next, we examined targets of the mTOR pathway. 4E-BP1, a direct target of mTORC1, showed an increase in phosphorylation upon CDKi, and accumulated in a hypophosphorylated form at 16 and 24 hours of Ro3306 treatment (Fig. 4A and 4B). Since 4E-BP1 phosphorylation controls the integrity of the cap-binding complex (Sonenberg and Hinnebusch, 2009), we carried out cap pulldown experiments using 7-methyl-GTP agarose beads. As expected, inhibition of mTORC1 using Torin1 (Thoreen et al., 2009) led to dissociation of eIF4G, eIF4A1 and eIF3B from eIF4E (Fig. S3C and S3D). Inhibition of CDK1 by treatment with Ro3306 for 4 or 16 hours, however, did not interfere with integrity of the cap-binding complex (Fig. S3C and S3D), indicating that CDKi does not repress translation via inhibition of mTOR signaling.

RPS6, a direct target of S6 kinase 1 (S6K1) and indirect target of mTORC1, was found to be strongly dephosphorylated early upon CDK1i (Fig. 4A and 4B). We first examined whether S6K1 mediates CDK1-dependent control of translation by generating HeLa cells that stably overexpress wild type (WT) or constitutively active (CA) S6K1. Phosphorylation levels of RPS6 were partially restored in the S6K1 overexpressing cells treated for 4 hours with Ro3306 (Fig. S4A), and translation suppression upon CDK1i was slightly, though significantly, reduced in comparison to control HeLa cells (Fig. 4D and S4B). We then tested whether RPS6 phosphorylation is responsible for this effect. In MEFs expressing bi-allelic phospho-deficient RPS6^P−/−^ (Ruvinsky et al., 2005), Ro3306 treatment suppressed translation to the same degree as in control RPS6^P+/+^ MEFs (Fig. 4E and S4C). Taken together, these results indicated that eIF2α phosphorylation and S6K1 activity contribute to translation control by CDK1, whereas RPS6 phosphorylation is not involved.

### CDK1 affects phosphorylation of translation-associated factors

CDK1 was recently detected in a ribosome interaction capture mass spectrometry analysis (Simsek et al., 2017), and found to phosphorylate ribosomal protein RPL12 (Imami et al., 2018). Together with our observation that CDK1i affects RPS6 phosphorylation (Fig. 4A and 4B), these findings prompted us to explore whether CDK1 might influence more generally the phosphorylation of ribosomal proteins and/or ribosome-associated factors. First, we explored if CDK1 indeed interacts with ribosomes. Polysome profile analysis revealed that a small proportion of CDK1 co-migrates with polysomes, and shifts to lighter fractions upon disassembly of polysomes by RNaseI (Fig. 5A and S5A).

**Figure 5.**
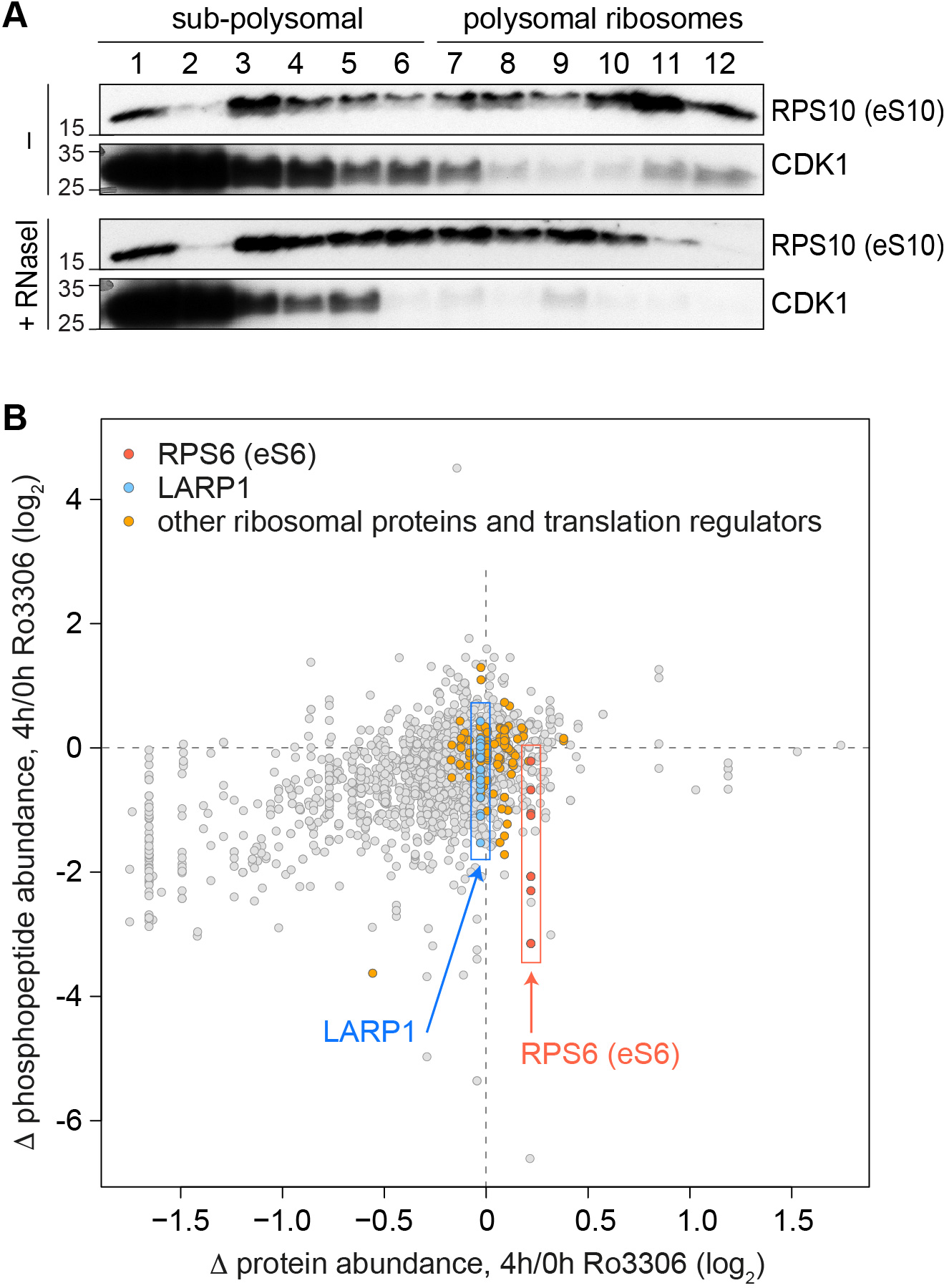
CDK1-dependent phosphorylation events associated with ribosomes. **(A)** HeLa cell lysates, either untreated or subjected to RNase I digestion, were fractionated following sucrose density gradient centrifugation. Association of RPS10 and CDK1 with the different fractions was monitored by Western blot analysis. **(B)** For phosphoproteomics of ribosomal fractions, HeLa cells were SILAC-labeled and either treated with DMSO or Ro3306 for 4 h. After lysis and disassembly of polysomes in low magnesium buffer, samples were mixed, and ribosomal fractions obtained by sucrose density centrifugation were subjected to phosphopeptide enrichment using PhosSelect iron affinity gel IMAC beads and analyzed by mass spectrometry followed by MaxQuant analysis. For all phosphopeptides detected under both conditions, the ratio (∆ phosphopeptide abundance, 4h/0h Ro3306) was plotted against the ratio of the corresponding total protein (∆ protein abundance, 4h/0h Ro3306). Phosphopeptides derived from LARP1 (blue), RPS6 (red) and other translation regulators (orange) are color-coded.

We then sought to identify possible targets of CDK1 associated with ribosomes using SILAC-based phosphoproteomics. Ribosomal fractions were obtained through sucrose gradient centrifugation from HeLa cells treated with either DMSO or Ro3306 for 4 hours, and subjected to mass spectrometry analysis. Phosphopeptide enrichment using PhosSelect iron affinity gel IMAC beads led to the identification of 2918 phosphorylated residues (Table S3). Ro3306-sensitive sites were detected in several ribosomal proteins (RPS6, RPS10, RPS17, RPL12, RPL29), translation factors (eIF2B4, eIF3 subunits, eIF4A1, eIF4B, eIF4GI, eIF5B), translation regulators (LARP1, YTHDF1) as well as mRNA splicing and export factors (SRRM2, SFSWAP, THOC2, ZC3H11A and eIF4A3). A prominent reduction of phosphorylation upon CDKi was observed for RPS6 and LARP1 (Fig. 5B), and this result could be verified in a second LC-MS/MS analysis using a smaller scale, TiO_2_-based enrichment of phosphopeptides (Fig. S5B and Table S4).

### CDK1 strongly enhances 5’TOP mRNA translation via LARP1

To gain further insight into the translation regulatory function of CDK1, we next performed ribosome footprint (Ribo-Seq) analysis. Cell cycle phase-dependent effects were avoided by using RPE1 cells arrested in G0 through serum starvation, and ribosome density (RD) was measured at an early time point (4 hours) after CDK1i. As an internal standard, equal amounts of a yeast lysate were spiked into the RPE1 cell lysates prior to RNaseI digestion, which allowed us to assess both the global and transcript-specific effects of Ro3306 on translation. To reduce distortion of results through ligation biases, the input RNA was fragmented by alkaline hydrolysis and subjected to the same library preparation protocol as the ribosomal footprints. Quality assessment showed the desired read lengths (Fig. S6A), pronounced periodicity and ORF enrichment for the footprints, but not the input RNA (Fig. S6B), as well as adequate reproducibility between biological replicates (Fig. S6A–C). As expected, CDK1i led to a global drop in RD (Fig. 6A, most transcripts below the diagonal), which corresponds to a two-fold reduction in the average RD (Fig. 6B). This result is in good agreement with the 3-fold reduction in polysomal ribosomes measured by polysome profiling (Fig. 3D).

**Figure 6.**
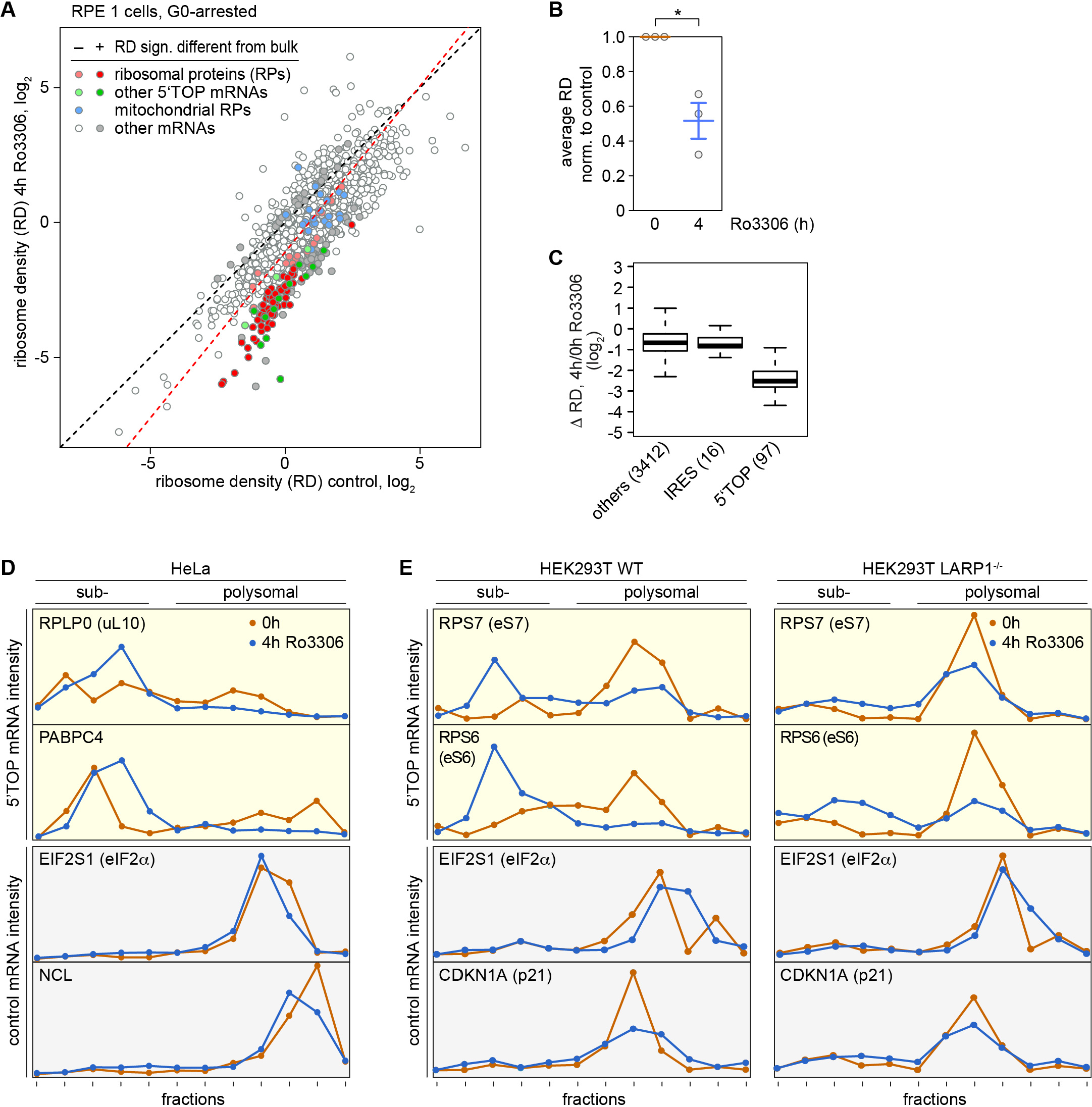
LARP1-dependent suppression of 5’TOP mRNA translation upon CDK1i. **(A)** For Ribo-Seq analysis, RPE1 cells were serum-starved for 48 h followed by a 4 h treatment with DMSO or Ro3306 (10 μM). Equal amounts of a yeast lysate was spiked into the DMSO- and Ro3306-treated samples. Ribosome densities (# ribosome footprints / # ORF-spanning reads in input RNA) were calculated after normalization to the yeast spike-in footprints from n = 3 biological repeat experiments. **(B)** Based on the Ribo-Seq analysis in (A), the average ribosome density was calculated after normalization to the yeast spike-in. Statistical significance was determined by one-sample Student’s t test; *, p ≤ 0.05. **(C)** Based on the Ribo-Seq analysis in (A), the fold-change in ribosome density (Δ RD) was calculated for IRES-containing mRNAs, 5’TOP mRNAs and all other mRNAs. **(D)** Polysome association of 5’TOP (RPLP0 and PABPC4) and ORF size-matched non-TOP (EIF2S1 and NCL) mRNAs was analyzed by polysome fractionation and subsequent qPCR analysis from DMSO- or Ro3306-treated (10 μM, 4 h) HeLa cells. **(E)** Polysome association of 5’TOP (RPS6 and RPS7) and ORF size-matched non-TOP (EIF2S1, CDKN1A) mRNAs was analyzed by polysome fractionation and subsequent qPCR analysis from DMSO- or Ro3306-treated (10 μM, 4 h) HEK293T WT or LARP1^−/−^ cells.

The analysis of individual transcripts revealed that CDK1i causes pronounced suppression of 5’TOP mRNAs, which includes all mRNAs encoding cytosolic ribosomal proteins (Fig. 6A and 6C). In contrast, mRNAs encoding mitochondrial ribosomal proteins, which do not contain a 5’TOP motif, or IRES-containing mRNAs, were not particularly sensitive to CDK1i (Fig. 6A and 6C). We confirmed in HeLa cells that Ro3306 treatment preferentially reduces the association of 5’TOP mRNAs (RPLP0 and PABPC4) with polysomes, whereas control mRNAs (EIF2α and NCL) were barely affected (Fig. 6D, repeats shown in Fig. S7A and S7B).

5’TOP mRNA translation was shown to be controlled by La related protein 1 (LARP1), which directly competes with eIF4E for binding to the cap of these transcripts (Lahr et al., 2017). Although LARP1 was initially reported to enhance translation of 5’TOP mRNAs under normal growth conditions (Tcherkezian et al., 2014), more recent evidence suggests that LARP1 actively represses 5’TOP mRNA translation (Fonseca et al., 2015; Philippe et al., 2018). This switch in activity appears to be controlled by phosphorylation of LARP1 (Hong et al., 2017). Since our phosphoproteomics analysis had indicated prominent changes in LARP1 phosphorylation (Fig. 5B), we decided to test whether the inhibitory effect of CDK1i on 5’TOP mRNA translation was dependent on LARP1 using knockout cells (doi.org/10.1101/491274). Whereas CDK1i led to a strong reduction of 5’TOP mRNA association with polysomes in WT HEK293T cells, the effect was much smaller in HEK293T LARP1^−/−^ cells (Fig. 6E, repeats shown in Fig. S7C and S7D). Since both HEK293T WT and LARP1^−/−^ cells responded to CDK1i by a reduction of polysomes (Fig. S7E–G), we concluded that LARP1 is not linked to the effect on global protein synthesis, while it does mediate the specific effect of CDK1 on enhancing 5’TOP mRNA translation.

## DISCUSSION

Early experiments measuring the incorporation of radiolabelled nucleosides and amino acids had already pointed to a tight connection between the proliferation rate and the rate of protein synthesis in cultured fibroblast subjected to contact inhibition (Levine et al., 1965) or serum deprivation (Rudland, 1974). Current concepts on mechanisms that couple the two rates focus on the mTOR signaling network, which integrates cues from growth factors and nutritional sensing in order to control a cell growth checkpoint in late G1 (Foster et al., 2010). The connection is based upon the notion that active mTOR, amongst its many effector functions, promotes cell proliferation as well as protein synthesis and ribosome biogenesis (Laplante and Sabatini, 2012).

Our SG-based screen for potential activators of translation revealed several candidate kinases that are primarily associated with cell cycle, proliferation and DNA damage (Fig. 1). A similar observation was made in an earlier screen by the Pelkmans lab, where inhibitors of several cell cycle kinases were found to prevent the dissolution of SGs (Wippich et al., 2013). These findings prompted us to test whether cell cycle kinases may be directly involved in controlling protein synthesis, and we decided to focus on CDK1 given its central role in driving the cell cycle (Santamaria et al., 2007).

Our analysis uncovered a novel, cell cycle-independent function of CDK1 in enhancing overall protein synthesis. We found that global protein synthesis rates are strongly reduced upon pharmacological inhibition of CDK1 using Ro3306 (Fig. 2) or Roscovitine (Fig. S1), and specificity of this observation could be confirmed by genetic inactivation of CDK1 in HT1080-derived HT2-19 cells (Fig. 2E). The effect was general as Ro3306 suppressed translation in transformed HeLa cells (Fig. 2) as well as in non-transformed RPE1 cells (Fig. 3D), HEK293T cells (Fig. S7E) and MEFs (Fig. S3A and S4C).

Previous studies identified several translation-associated factors as direct substrates of CDK1 in mitosis, including S6K1 (Papst et al., 1998; Shah et al., 2003), Raptor (Ramirez-Valle et al., 2010), eEF2K (Smith and Proud, 2008), eIF4GI (Dobrikov et al., 2014), 4E-BP1 (Heesom et al., 2001; Shuda et al., 2015; Velasquez et al., 2016), eEF1D (Monnier et al., 2001; Sivan et al., 2011) and the ribosomal protein RPL12 (Imami et al., 2018). These studies suggested that CDK1 regulates translation only during mitosis, and many of the reported CDK1-dependent phosphorylation events (S6K1, eIF4GI, eEF1D) were proposed to repress global translation in order to increase the translation of mitosis-specific transcripts. On the other hand, CDK1-dependent phosphorylation of 4E-BP1 (Heesom et al., 2001; Shuda et al., 2015) and eEF2K (Smith and Proud, 2008) were linked to a positive role of CDK1 in global translation during mitosis. Our findings suggest that CDK1 also exerts a translation regulatory function independently of the cell cycle phase, since Ro3306 treatment suppressed translation in cells arrested in early S-phase (Fig. 3C) or in G0 (Fig. 3D). Thus, we propose that in addition to controlling translation during mitosis, CDK1 serves as a relay to balance the overall proliferation rate of a cell with the overall protein synthesis rate. This may be linked to the notion that CDK1 phosphorylates different substrates depending on its activity during the cell cycle, its subcellular localization, its association with co-activators (cyclins or RINGO proteins) and possibly its phosphorylation status (Gupta et al., 2007; Hochegger et al., 2008; Nebreda, 2006; Swaffer et al., 2016)

When addressing the mechanism by which CDK1 enhances global protein synthesis, we found that CDK1 influences translation initiation via multiple, possibly redundant pathways. First, we found an increase in eIF2α phosphorylation upon CDKi (Fig. 4A), and since translation suppression was reduced in MEFs expressing non-phosphorylatable eIF2α S51A (AA) (Fig. 4C), one role of CDK1 is to promote recharging of the eIF2-GTP-tRNA_i_^Met^ ternary complex. Second, we observed a pronounced reduction in RPS6 phosphorylation upon CDKi (Fig. 4A and 5B), and reduced translation suppression in HeLa cells overexpressing S6K1 (Fig. 4D) indicated that CDK1 also enhances translation initiation through the S6K1 signaling axis. Of note, CDK1 most likely does not act via mTOR since CDK1i, in contrast to mTOR inhibition, did not alter the integrity of the cap binding complex (Fig. S3C and S3D). Third, we could show that CDK1 co-sediments with polysomes (Fig. 5A), and a recent mass spectrometry approach found CDK1 to be associated with ribosomes (Simsek et al., 2017). This is in line with the notion that RPL12 is a known substrate of CDK1, and RPL12 phosphorylation was recently shown to enhance a mitotic translation program (Imami et al., 2018). Hence, it is possible that CDK1 stimulates global translation by phosphorylating additional proteins of, or associated with, the ribosome. Finally, our results show that CDK1 is a pronounced activator of 5’TOP mRNA translation, which includes the synthesis of all ribosomal proteins (Fig. 6). Hence, CDK1 has a sustained effect on global protein synthesis in proliferating cells as it enhances translation at the initiation level, possibly also at the elongation level (Smith and Proud, 2008), and by promoting biogenesis of the protein synthesis machinery.

We were intrigued by the pronounced effect of CDK1i on 5’TOP mRNA translation. In agreement with this finding, it is well known that cell cycle progression tightly correlates with 5’TOP mRNA translation. For example, cell cycle arrest in G0, at the beginning of S-phase or in M-phase, strongly reduces translation of 5’TOP mRNAs (Meyuhas and Kahan, 2015). Likewise, translation of 5’TOP mRNAs is low in resting adult liver cells, but high in developing fetal liver cells as well as in proliferating adult liver cells during regeneration (Aloni et al., 1992). We propose that CDK1 has a central role in coupling 5’TOP mRNA translation with the proliferation status of the cell since i) LARP1 phosphorylation is strongly dependent on CDK1 activity (Fig. 5B) and ii) CDK1 controls 5’TOP mRNA translation in a LARP1-dependent manner (Fig. 6E). Future studies will need to show if LARP1 is a direct target of CDK1, and address the detailed mechanism by which CDK1 antagonizes the inhibitory activity of LARP1 on 5’TOP mRNA translation.

Taken together, our results suggest that CDK1 acts as a central relay connecting proliferative cues with protein synthesis. This activity occurs in parallel to the mTOR kinase, which functions as a signaling hub that couples cues from growth factors and nutrient sensing with protein synthesis. CDK1 and mTOR thereby share common targets including S6K1, 4EBP1 and LARP1. Together with mTOR and Ras/Erk, CDK1 appears to form a homeostatic network that coordinates proliferative cues and growth signals with the availability of the protein synthesis machinery and the rate of protein synthesis.

## Acknowledgments

We would like to thank Andy Porter (Imperial College Faculty of Medicine, London, UK) for generously providing the HT1080 and HT2-19 cell lines, Randal Kaufman (Sanford Burnham Prebys Medical Discovery Institute, La Jolla, CA, USA) for the eIF2α-AA and -SS MEFs, and Oded Meyuhas (Hebrew University of Jerusalem, Israel) for the RPS6^P+/+^ and RPS6^P−/−^ MEFs. We also thank John Blenis (Weill Cornell Medicine, New York, NY, USA) for the S6K1 plasmids, Jamal Tazi (Institut de Génétique Moléculaire de Montpellier, France) for the G3BP1 cDNA, Atsushi Miyawaki (RIKEN Center for Brain Science, Japan) for the FUCCI plasmids, and Thomas Mayo (Brigham and Women’s Hospital, Harvard Medical School, Boston, MA, USA) for help with cloning GFP-G3BP1. We are thankful to Ulrike Friedrich, Guenter Kramer and Bernd Bukau (all at the Center for Molecular Biology of Heidelberg University, Germany) for providing yeast lysates and assistance with the Ribo-Seq protocol.

Mass spectrometry analysis was carried out by the Core Facility for Mass Spectrometry & Proteomics at the ZMBH. FACS sorting was done by the FACS core facility at the ZMBH. siRNA library preparation was performed by the Cellnetworks Advanced Biological Screening Facility of Heidelberg University, and next generation sequencing was carried out by the Cellnetworks Deep Sequencing Core Facility of Heidelberg University.

This work was supported by grant SFB 1036 from the Deutsche Forschungsgemeinschaft (DFG) to G. Stoecklin and M. Knop; grant TRR 186 from the DFG to A. Ruggieri and G. Stoecklin; grant INST 35/1067-1 FUGG to M. Knop; and grant 6108-00197B from The Danish Council for Independent Research - Natural Sciences to C.K. Damgaard.

## Author contributions

K. Haneke performed the screen and carried out most experiments, J. Schott did the Ribo-Seq bioinformatics analysis, D. Lindner prepared the Ribo-Seq libraries, C. Mongis and M. Knop were instrumental for the imaging screen, A. Ruggieri generated HeLa-FUCCI cells, A.K. Hollensen and C.K. Damgaard generated the LARP1^−/−^ HEK293T cells, K. Haneke and G. Stoecklin designed the study, analyzed the data and wrote the manuscript; and all authors approved the manuscript.

## MATERIALS and METHODS

### Plasmid generation

The GFP-G3BP1 sequence was obtained from J. Tazi (Institut de Génétique Moléculaire de Montpellier, France) and cloned into the NheI and EcoRI sites of pCI-puro, resulting in pCI-puro-GFP-G3BP1 (p2163). pKH3-HA-S6K1-WT (p2760) and pKH3-HA-S6K1-CA (F5A-T389E-R5A) (p2762) were kindly provided by J. Blenis (Weill Cornell Medicine, New York, NY, USA). HA-S6K1-WT and HA-S6K1-CA sequences were amplified using oligos G4542 and G4543 from plasmid p2760 and p2762, respectively, and cloned into the SmaI sites of pWPI-BLR (Ruggieri et al., 2012), resulting in the generation of pWPI-BLR-HA-S6K1-WT (p3669) and pWPI-BLR-HA-S6K1-CA (F5A-T389E-R5A) (p3671). pWPI-FUCCI-Kusabira-Orange-Cdt1-Zeo and pWPI-FUCCI-mVenus-Geminin-Zeo were generated by EcoRI-XbaI excision of the human Cdt1 gene N-terminally fused to mKO2 from plasmids pCSII-EF-MCS-mKO2-hCdt1-(30/120) and of the human Geminin gene N-terminally fused to mVenus from pCSII-EF-MCS-mVenus-hGeminin-(1/110), respectively (both kindly provided by A. Miyawaki, RIKEN Center for Brain Science, Japan) (Sakaue-Sawano et al., 2008). Both sequences were inserted into the lentiviral transduction vector pWPI carrying a zeocin resistance gene.

### Generation of stable cell lines and knockout cell lines

Hela GFP-G3BP1 cells were generated by plasmid transfection of pCI-puro-GFP-G3BP1 (p2163) using PEI. 24 h after transfection, cells were subjected to selection pressure by the addition of 2 μg/ml puromycin (Gibco). After two weeks of selection, mass cell cultures were FACS-sorted using a BD FACSAria IIIu cell sorter. HeLa-FUCCI-Kusabira-Orange-hCdt1 and HeLa-FUCCI-mVenus-hGeminin cells were generated by lentiviral transduction of pWPI-FUCCI-Kusabira-Orange-Cdt1-Zeo and pWPI-FUCCI-mVenus-Geminin-Zeo, respectively. Retroviral transduction and generation of stable cell lines was performed as described in (Ruggieri et al., 2012). In short, 293T cells were seeded into 6 cm-diameter dishes and transfected using the CalPhos mammalian transfection kit (Becton Dickinson) as recommended by the manufacturer. For transfection, the packaging plasmid (pCMV∆8.91), the transfer vector (pWPI-based) and the VSV envelope glycoprotein expression vector (pMD2.G) were used in a concentration ratio of 3:3:1. Transduction of HeLa cells with the lentiviral particles was repeated three times every 12 h to achieve high number of integrates and thus high expression levels. Transduced cell pools were subjected to selection with medium containing 100 μg/ml zeocin (Invitrogen) and high expressing cells were sorted by FACS. HeLa-HA-S6K1-WT and HeLa-HA-S6K1-CA cells were generated similarly by retroviral transduction using pWPI-BLR-HA-S6K1-WT (p3669) and pWPI-BLR-HA-S6K1-CA (F5A-T389E-R5A) (p3671). Cells were subjected to selection pressure by the addition of 5 μg/ml blasticidin (Invitrogen). To create plasmids for expression of LARP1-specific gRNAs, LARP1-oligo1 and LARP1-oligo2 were annealed and cloned into Esp3I-digested LentiCRISPRv2, resulting in a vector designated LentiCRISPR-LARP1gRNA. HEK293T cells were maintained in DMEM medium (Gibco) supplemented with 10% fetal calf serum (Gibco) and 100 units/mL penicillin/streptomycin (Gibco). All cells were cultured at 37°C in 5% (v/v) CO2. One day prior to transfection, HEK293T cells were seeded at a density of 3 x 105 cells/well in 6-well plates. Transfections were carried out using 1 μg LentiCRISPR-LARP1 gRNA and Lipofectamine 2000 Transfection Reagent (Invitrogen) according to manufacturer’s protocol. Two days after transfection, the cells were reseeded at a density of 0.2 cells/well in 96-well plates. After expansion of single cells, genomic DNA was purified using the GenElute Mammalian Genomic DNA Miniprep Kits (Millipore) according to manufacturer’s protocol. LARP1 CRISPR/Cas9 KO was verified by PCR on genomic DNA using the primers LARP1-oligo3 and LARP1-oligo4, followed by Sanger sequencing of the resulting PCR-product using the primer LARP1-oligo3.

### Cell culture

HeLa cells, HEK293T cells and MEFs were maintained in Dulbeccos’s modified Eagle’s medium (DMEM, Gibco) containing 10% fetal calf serum (FCS, PAA Laboratories), 2 mM L-glutamine, 100 U/ml penicillin and 100 μg/ml streptomycin (all PAN Biotech). eIF2α Ser51 SS (WT) and AA MEFs (Scheuner et al., 2001) were a kind gift from R. Kaufmann (Sanford Burnham Prebys Medical Discovery Institute, La Jolla, CA, USA); RPS6^P+/+^ and RPS6^P−/−^ MEFs were generously provided by O. Meyuhas (Hebrew University of Jerusalem, Israel). The generation of HEK293T LARP1^−/−^ cells will be published elsewhere (doi.org/10.1101/491274). HT1080 and HT2-19 cells (Itzhaki et al., 1997) were a kind gift from A. Porter (Imperial College School of Medicine, London, UK) and were maintained in Dulbeccos’s modified Eagle’s (DMEM, Gibco) high glucose and non-essential amino acids (NEAA) medium containing 10% fetal calf serum (FCS, PAA Laboratories), 2 mM L-glutamine, 10 mM pyruvate, 40 U/ml penicillin and 40 μg/ml streptomycin (all PAN Biotech). HT2-19 cells were additionally supplemented with 0.2 mM IPTG (AppliChem). For CDK1 depletion HT2-19 cells were seeded at very low density and cultured in the absence of IPTG for 7 days. RPE1 cells, kindly provided by I. Hoffmann (German Cancer Research Center, Heidelberg), were cultured in HAM’s F-12 medium (HAM’s F-12 (1:1), Millipore) containing 10% FCS, 2 mM L-glutamine, 100 U/ml penicillin and 100 μg/ml streptomycin. HeLa-FUCCI-Kusabira-Orange-hCdt1, HeLa-FUCCI-mVenus-hGeminin and HeLa-GFP-G3BP1 cells were FACS sorted, cultured without selection pressure and maintained at low passage numbers. All cells were cultured at sub-confluency, at 37°C in 5% CO_2_. For treatment with inhibitors, cells were seeded the evening before, and Ro-3306 (Sigma, 10 μM), Roscovitine (Sigma, 20 μM), Torin-1 (200 nM, Tocris Bioscience) or control solvent (DMSO) were diluted in fresh medium, which was added onto the cells for the indicated times. For synchronization, HeLa cells were subjected to a double thymidine block following standard procedures (18 h 2 mM thymidine, 9 h release, and 18 h 2 mM thymidine).

### Screening approach and SG score

For the siRNA screen, 96-well MGB096-1-2-LGL matriplates (Brooks) were coated with a siRNA transfection mix containing the Dharmacon siGenome siRNA libraries GU-003505 Human Protein kinase and GU-003705 Human Phosphatase from Thermo Scientific, Lipofectamine RNAiMAX (Invitrogen), 57 mM sucrose, 0.03% gelatine/fibronectin in solution and OPTIMEM. The coated plates were prepared for long term storage by drying, coating and drying were carried out in the Cellnetworks Advanced Biological Screening Facility of Heidelberg University using the Hamilton “STAR” pipetting robot. The siRNA libraries were directed against in total 711 human kinases and 256 human phosphatases including 4 individual siRNAs per gene. 2000 HeLa-GFP-G3BP1 cells per well were seeded into the 96-well plates, siRNAs were transfected at a final concentration of 50 nM and kd was carried out for 72 h. Cells were fixed for 10 minutes at RT using 4% PFA in PBS supplemented with Hoechst dye (1:10000 diluted). Afterwards, cells were washed 3 times and stored in PBS at 4°C and in the dark until examination under the microscope. Seeding, washing and fixation were done with a microplate suspensor (Thermo Scientific Multidrop Combi) in order to ensure fast, synchronous and equal handling. SG formation was analyzed using a Nikon eclipse Ti-E microscope and a Nikon plan Apo 60x oil objective that was constantly supplied with immersion oil by a pumping system. 16 images per well were taken automatically using a sCMOS camera (Flash4, Hamamatsu), Nikon JOBS software and the Nikon perfect focus system, and images were subsequently analyzed by eye. For every phosphotransferase, a SG score was calculated by multiplying the sum of SG-containing cells, observed with all 4 siRNAs, by the number of siRNAs causing SGs.

### Immunofluorescence (IF) and microscopy

Cells were seeded onto glass coverslips one day before drug treatment. Cells were fixed with 4% paraformaldehyde (PFA) for 10 min, permeabilized with 0.5% Triton-X in PBS for 10 min and blocked with 3% BSA in PBS for 1 h at RT. Cy3- or Cy2-conjugated secondary donkey antibodies (Jackson ImmunoResearch Laboratories, West Grove, PA) were used for detection of primary antibodies. DNA was stained with Hoechst dye (1: 10000, Sigma). Coverslips were mounted onto glass slides using a solution of 14% polyvinol-alcohol (P8136, Sigma) and 30% glycerol in PBS. Microscopy was performed on a Leica DM 5000 Microscope using a 20x or 40x dry objective, or a 40x oil objective. Alternatively, a Nikon eclipse Ti-E microscope was used in combination with a 40x dry objective or a 60x oil objective. Images were taken with an Andor CCD camera or a pco edge sCMOS camera, and subsequently processed and analyzed using Adobe Photoshop and Fiji software.

### Western blot analysis

Cells were lysed by scraping in ice-cold protein lysis buffer (50 mM Tris-HCl pH 7.4, 150 mM NaCl, 15 mM MgCl_2_, 1% Triton X-100) supplemented with EDTA-free protease inhibitor cocktail (Roche) and phosphatase inhibitors (1 mM sodium vanadate, 50 mM sodium fluoride, 0.04 μM okadaic acid). Samples were incubated for 5 min on ice and nuclei were removed by centrifugation for 5 min at 10,000 g at 4°C. 10–20 μg total protein was diluted in SDS sample buffer (4 % SDS, 20 % Glycerol, 10 % DTT, 0.004 % Bromphenol Blue, 0.125 M Tris HCl), loaded onto 5–20% polyacrylamide gradient gels and transferred to a 0.2 μm pore size nitrocellulose membrane (PeqLab) by wet blotting. Membranes were blocked in 5% milk or 5% BSA (both diluted in PBS) at RT, incubated with primary antibodies diluted in PBSA overnight at 4°C and washed with TBS containing 1% Tween 20 (TBST). Horseradish peroxidase (HRP)-conjugated secondary antibodies (Jackson Immunoresearch, diluted 1:5000 in PBS) and Western Lightning Enhanced Chemiluminescence substrate (Perkin Elmer) were used for detection.

### Antibodies

Mouse anti-G3BP1 (TT-Y) (Santa Cruz sc-81940), mouse anti-acetylated tubulin (Sigma C3B9), goat anti-eIF3B (Santa Cruz sc-16377), mouse anti-puromycin (Millipore MABE343), mouse anti-CDK1 (B-6) (Santa Cruz sc-8395), rabbit anti-RPS6 (5G10) (Cell Signaling #2217), rabbit anti-phospho-eIF2alpha (Cell Signaling #9721), rabbit anti-eIF2alpha (Cell Signaling #9722), rabbit anti-4E-BP1 (Cell Signaling #9644), rabbit anti-phospho-4E-BP1 (Thr37/46) (Cell Signaling #236B4), rabbit anti-phospho-RPS6 (Ser235/236) (D57.2.2E) (Cell Signaling #4858), mouse anti-tubulin (DM1A) (Sigma T9026), mouse anti-HA.11 (MMS-101P, Covance), rabbit anti-RPS10 (Abcam ab151550), mouse anti-RPS3 (Santa Cruz sc-376098), rabbit anti-LARP1 (Abcam ab86359), mouse anti-FLAG M2 (Sigma F 3165), mouse anti-eIF4E (P-2) (Santa Cruz sc-9976), rabbit anti-eIF4G (Santa Cruz sc-11373), goat anti-eIF4AI (Santa Cruz sc-14211)

### Primers

G1714, 5’-GAAGGCTCATGGCAAGAAGG-3’ (beta globin fw)

G1715, 5’-ATGATGAGACAGCACAATAACCAG-3’ (beta globin rev)

G2943 5’-TGGAGACTCTCAGGGTCGAAA-3’ (CDKN1A fw)

G2944 5’-GGCGTTTGGAGTGGTAGAAATC-3’ (CDKN1A rev)

G2979, 5’-TCGATGGGCGATCTATTTCCCTGT-3’ (NCL fw)

G2980, 5’-TGTTGCACTGTAGGAGAGGTTGCT-3’ (NCL rev)

G3007, 5’-GAGTTCGAGTCCGGCATCT-3’ (RPS7 fw)

G3008, 5’-CGACCACCACCAACTTCAA-3’ (RPS7 rev)

G4542: 5’-GATCCCCCGGGAATAACATCCACTTTGCCTTTCTC-3’

G4543: 5’-TTTCCCGGGTCATAGATTCATACGCAGGTGC-3’

G4737, 5’-TCTACAGAAAACATGCCCATTAAG-3’ (EIF2S1 fw)

G4738, 5’-GCCATAGCTTGACTGAGGACA-3’ (EIF2S1 rev)

G4739, 5’-TCTACAACCCTGAAGTGCTTGAT-3’ (RPLP0 fw)

G4740, 5’-CAATCTGCAGACAGACACTGG-3’ (RPLP0 rev)

G4753, 5’-GTAGGCCGTGCACAAAAGA-3’ (PABPC4 fw)

G4754, 5’-AATGTAGAGATTCACCCCCTGA-3’ (PABPC4 rev)

G4976, 5’-CTGGGTGAAGAATGGAAGGGTT-3’ (RPS6 fw)

G4988, 5’-TGCATCCACAATGCAACCAC-3’ (RPS6 rev)

LARP1-oligo1, 5’-CACCGAGACACATACCTGCCAATCG-3’

LARP1-oligo2, 5’-AAACCGATTGGCAGGTATGTGTCTC-3’

LARP1-oligo3, 5’-GGGAAAGGGATCTGCCCAAG-3’

LARP1-oligo4, 5’-CACCAGCCCCATCACTCTTC-3’

### Polysome profile analysis

Cells were seeded one day before the experiment and kept at sub-confluency in order to prevent translation suppression by contact inhibition. Cells were then treated with 100 μg/ml cycloheximide (CHX) for 5 min at RT in order to stabilize existing polysomes before washing with ice-cold PBS and harvesting by scraping in polysome lysis buffer (20 mM Tris HCl pH 7.5, 150 mM NaCl, 5 mM MgCl_2_, 1 mM DTT, 100 mg/ml CHX, 1% Triton X-100, 40 U/ml RNasin, EDTA-free complete protease inhibitors (Roche)). Lysates were rotated end over end for 10 min at 4°C and cleared by centrifugation at 10,000 g for 10 min at 4°C. 40 μl lysate were saved for Western blot analysis before the cellular lysate was loaded onto linear 17.5–50% sucrose gradients (dissolved in 20 mM Tris-HCl pH 7.5, 5 mM MgCl_2_, 150 mM NaCl). Sucrose density gradient centrifugation was carried out at 35,000 rpm at 4°C using a SW60 rotor (Beckman) for 2.5 h. Polysome profiles were recorded by measuring the absorbance at 254 nm using a Teledyne ISCO Foxy Jr. or a Teledyne ISCO Foxy R1 system in combination with PeakTrak software. Profiles were then aligned manually according to the 80S peak, and the percentage of polysomal ribosomes was calculated by dividing the area under the curve of the polysomal ribosomes by the total area under the curve.

### Polysome fractionation

During gradient elution, fractions of approximately 300 μl were collected every 14 seconds. For RNA isolation, 300 μl Urea buffer (10 mM Tris pH 7.5, 350 mM NaCl, 10 mM EDTA, 1 % SDS and 7 M urea) containing 25 fmol rabbit HBB2 *in vitro* transcript and 300 μl Phenol:Chloroform:Isamylalcohol (PCI) (25:24:1) were added to each fraction. After phase separation, RNA was isolated from the aqueous phase and precipitated using isopropanol. RNA levels in the different fractions were subsequently analyzed by qPCR as follows: RNA was reverse transcribed using the MMLV reverse transcriptase (Promega), followed by cDNA amplification using the PowerUp SYBR Green Master Mix (Thermo Fisher Scientific) and the QuantStudio 5 Real-TimePCR system (Thermo Fisher Scientific). All CT values were normalized to the HBB2 spike-in transcript in order to correct for isolation differences.

For protein purification, 300 μl Tris-HCl (20 mM, pH 7.5) and 10 μl StrataClear beads were added to each fraction. Samples were rotated end over end at 4°C overnight, centrifuged at ~ 100 g for 2 min., and proteins were eluted from the beads using SDS sample buffer.

### Ribosome footprint (Ribo-Seq) analysis

RPE1 cells were cultured in the absence of FBS for 48 h. Afterwards, cells were incubated for 4 h in fresh medium without FBS supplemented with either DMSO or Ro3306, washed once in ice-cold PBS supplemented with 100 μg/ml CHX and harvested by scraping in polysome lysis buffer. Lysates were rotated end over end for 10 min at 4°C and cleared by centrifugation at 10,000 g for 10 min at 4°C. The DMSO- and Ro3306-treated samples were adjusted to the same OD260 before yeast polysome lysate (2% of the RPE1 lysates according to OD260 measurement) was spiked into each sample. 10% of the lysates were saved as input samples. The lysates were subsequently digested with RNase I (60 units per OD260) for 5 min at 4°C, and the reaction was stopped by addition of Superase Inhibitor (6 units). Samples were then fractionated by 17.5–50% sucrose density gradient centrifugation, and RNA was purified from the cytoplasmic lysate (input) or from the monosomal fractions (ribosome protected fragments) using PCI (25:24:1) by phase separation. Both input and ribosome protected fragments were depleted of rRNA with the Ribo-Zero Gold Kit (Illumina). Input RNA was randomly fragmented by alkaline hydrolysis at pH 10 for 12 min at 95°C. Fragmented RNA and ribosome protected fragments were size-selected (25 - 35 nt) on a 15% polyacrylamide TBE-urea gel. After end-repair with T4 PNK, 3 ng per sample were used for library preparation using the NEXTflex Small RNA-Seq Kit v3 according to the manufacturer’s manual. Libraries were multiplexed and sequenced on one lane of a NextSeq500 sequencer (Illumina).

For Ribo-Seq data analysis, adapter sequences were first removed with the FASTX-toolkit (http://hannonlab.cshl.edu/fastx_toolkit/), and the four random nucleotides at the beginning and end of the reads were trimmed. Read alignment was then performed using bowtie (Langmead et al., 2009). Reads that did not map to human tRNA or rRNA sequences were aligned to a common human transcriptome reference (wgEncodeGencodeBasicV27) and a yeast transcriptome (sacCer3ensGene). In order to summarize reads at the gene level, only reads that map to the annotated ORF of isoforms of one specific gene (as defined by a common gene symbol) were counted with an in-house-developed perl script. In order to identify individually regulated mRNAs with DESeq2, human read counts were normalized with the median ratio method before calculating average fold-changes, and p-values for changes in ribosome density were obtained from a likelihood ratio test (Love et al., 2014). The sum of read counts assigned to yeast or human ORFs was used in order to measure global changes in translation efficiency. For categorization, mRNAs that contain an IRES-element (according to http://iresite.org/IRESite_web.php?page=browse_cellular_transcripts) or a 5’TOP motif (according to (Meyuhas and Kahan, 2015)) were grouped.

### Puromycin incorporation

Cell were treated with 10 μg/ml puromycin (Gibco, Life Technologies) for 5 min at 37°C, washed twice with PBS and lysed in protein lysis buffer. For Western blot analysis, equal amounts of total cell lysates were separated by SDS-PAGE. Puromycin signals were detected with anti-puromycin antibody. The signal intensity was measured along the entire lane and normalized to the overall Ponceau S staining of the corresponding lane.

### Phosphoproteomics

HeLa cells were cultured in SILAC medium (DMEM without arginine, lysine, glutamine and pyruvate, containing 10% FBS for SILAC (Silantes), 2 mM L-glutamine, 100 U/ml penicillin and 100 μg/ml streptomycin (all PAN Biotech) and either light- or heavy-labeled amino acids (SILAC amino acids (Silantes 211603902 and 201603902)) for 14 days. 4 times 3.5 × 10^6^ heavy-labeled cells and 4 times 3.5 × 10^6^ light-labeled cells were seeded into in total 8 15-cm dishes. Light-labeled cells were treated with DMSO and heavy-labeled cells with Ro3306 (10 μM) for 4 h, and labels were swapped in the repeat experiment. Cells were washed with ice-cold PBS, 200 μl low magnesium polysome lysis buffer (20 mM Tris HCl pH 7.5, 150 mM NaCl, 0.25 mM MgCl_2_, 1 mM DTT, 100 mg/ml CHX, 1% Triton X-100, 40 U/ml RNasin, EDTA-free complete protease inhibitors (Roche), phosphatase inhibitors (PhosphoSTOP, Roche)) were added and lysates were harvested by scraping. Lysates were then rotated end over end for 10 min at 4°C and cleared by centrifugation at 10,000 g for 10 min at 4°C. The total protein content of the lysate was measured using a Bradford assay. Equal amounts of total protein from the heavy and the light sample were mixed and loaded onto low magnesium 17.5–50% sucrose gradients (dissolved in 20 mM Tris-HCl pH 7.5, 5 mM MgCl_2_, 150 mM NaCl). Sucrose density gradient centrifugation was carried out as described above, and ribosomal fractions were pooled. Proteins were precipitated using the Wessel-Flügge precipitation protocol (Wessel and Flugge, 1984). Samples were enriched for phosphopeptides using PhosSelect iron affinity gel IMAC beads (first repeat) or by TiO_2_-SIMAC-HILIC (TiSH) phosphopeptide enrichment and fractionation (second repeat), and subsequently subjected to mass spectrometry and Maxquant analysis at the Core Facility for Mass Spectrometry & Proteomics of the ZMBH. Go annotations were added using Perseus software.

### Cap pulldown assay

Cells were lysed by scraping in cap pulldown lysis buffer (50 mM Tris-HCl pH 7.8, 150 mM NaCl, 1 mM EDTA, EDTA-free complete protease inhibitors (Roche)). Lysates were then rotated end over end for 10 min at 4°C and nuclei were pelleted by centrifugation at 10,000 g for 10 min at 4°C. 10% of total cell lysates were saved as input samples. 50 μl of γ-aminophenyl-m^7^GTP agarose C10-linked beads (Jena Biosciences) were added to the remaining sample, which was then rotated end over end at 4°C overnight. Beads were washed 5 times using cap pulldown wash buffer (10 mM Tris-HCl pH 7.8, 150 mM NaCl, 0.1% NP40) and eluted with SDS sample buffer.

### Statistical analysis

Statistical analysis was performed using Microsoft Excel 2010 or Graph-Pad Prism software (GraphPad). Statistical significance was calculated by performing a one-tailed Student’s t test or a one-tailed one-sample t test (***, p < 0.001; **, p < 0.01; *, p < 0.05).

