## Supplemental Figures for "CDK1 couples proliferation with protein synthesis"

*Figure S1, Haneke et al.*

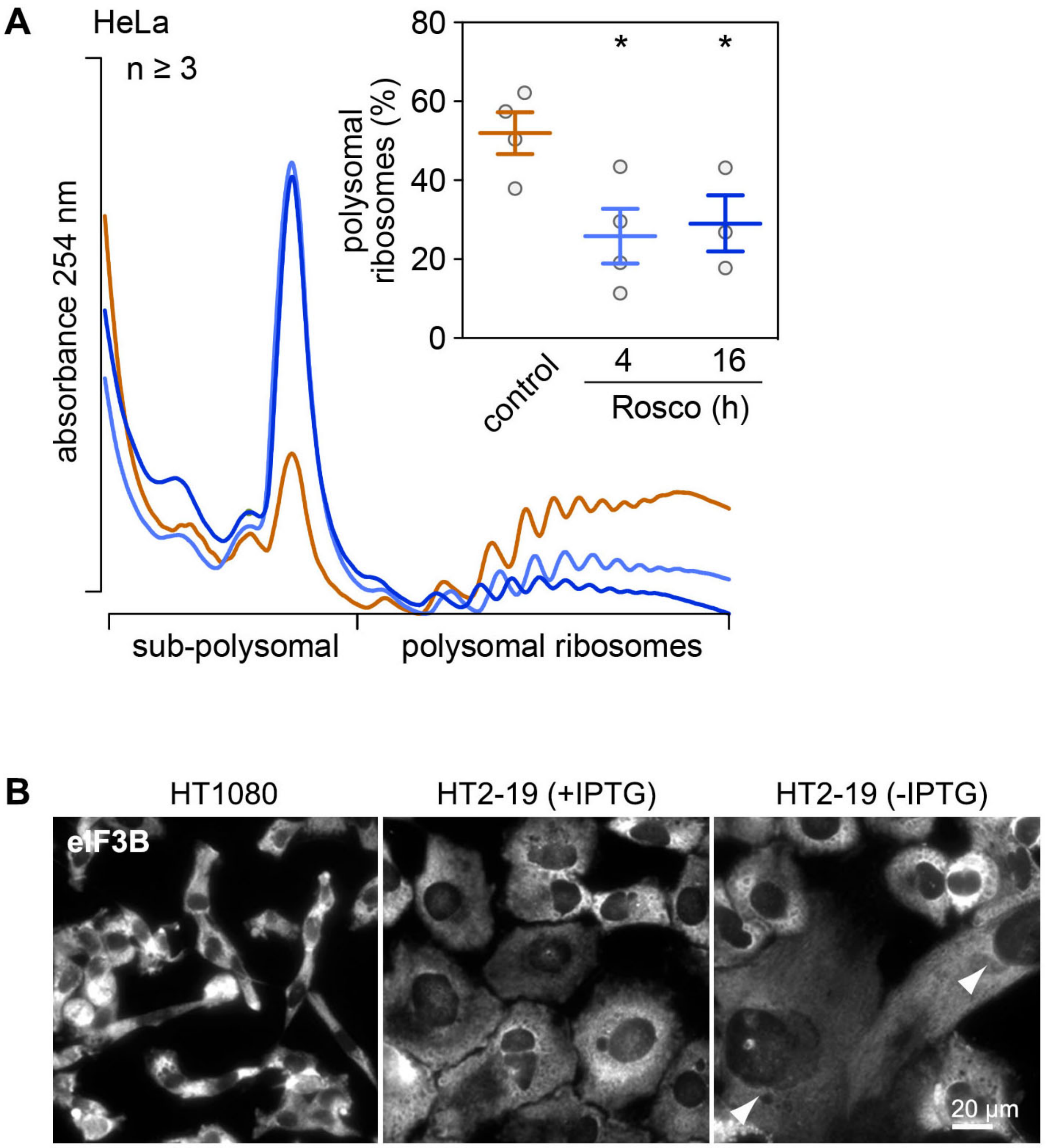


Figure S1. **Pharmacological inhibition and genetic depletion of CDK1. (A)** HeLa cells were treated either with control solvent (DMSO) or with the CDK1 inhibitor Roscovitine (20 µM) for 4 and 16 h. Polysome profiles were recorded after sucrose density gradient centrifugation and the percentages of polysomal ribosomes were calculated (average ± SEM, n ≥ 3). Statistical significance was determined by unpaired Student's t test; *, p ≤ 0.05. **(B)** HT1080 and HT2-19 cells were cultured in the presence or absence of IPTG (0.2 mM) for 7 days. Cellular morphology was assessed by IF microscopy of fixed cells stained with anti-eIF3B antibody.

*Figure S2, Haneke et al.*

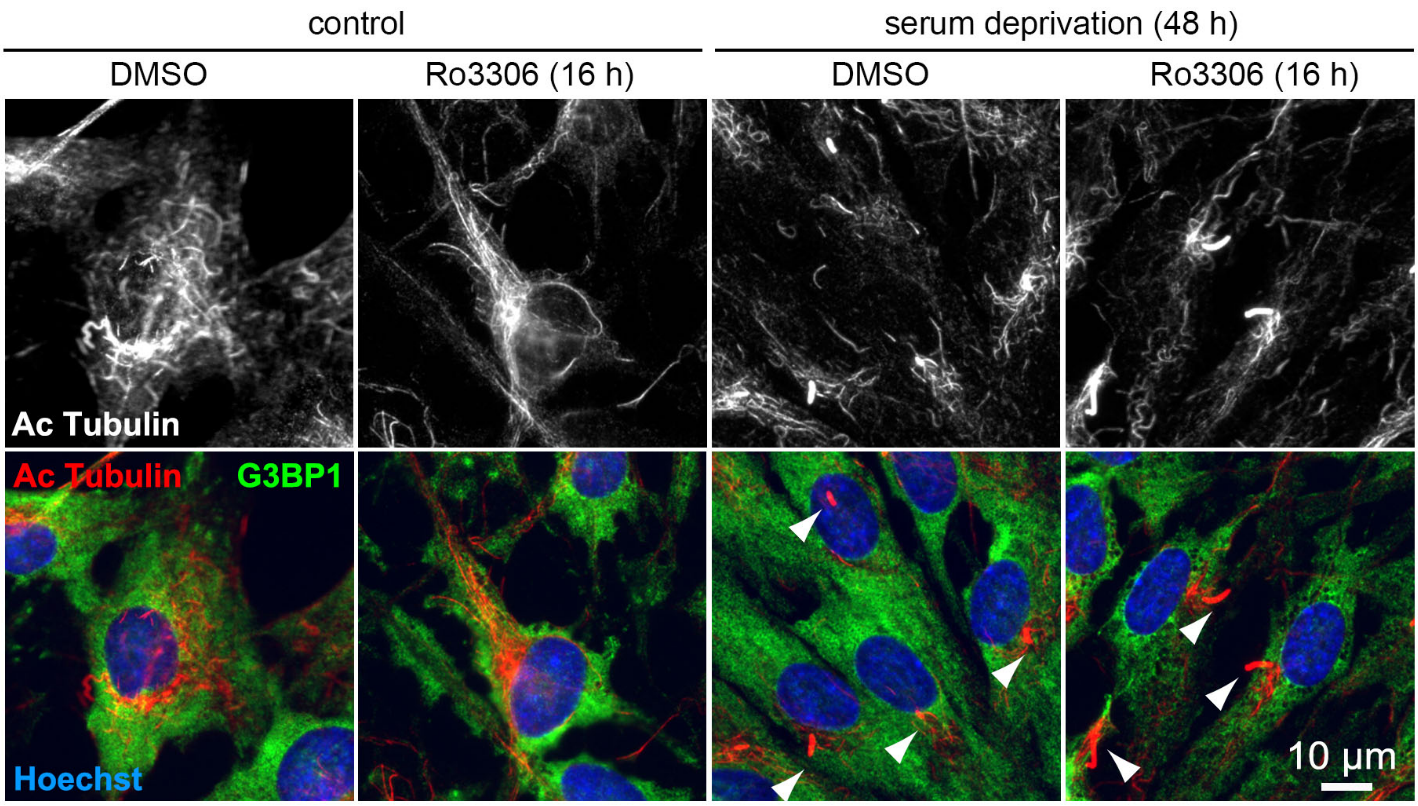


Figure S2. **Serum deprivation-induced cell cycle arrest in RPE1 cells.** RPE-1 cells were serum deprived for 48 h and subsequently treated with DMSO or Ro3306 (10 µM) for 16 h. Formation of cilia was analyzed by IF microscopy of fixed cells stained with anti-acetylated tubulin and anti-G3BP1 antibody.

*Figure S3, Haneke et al.*

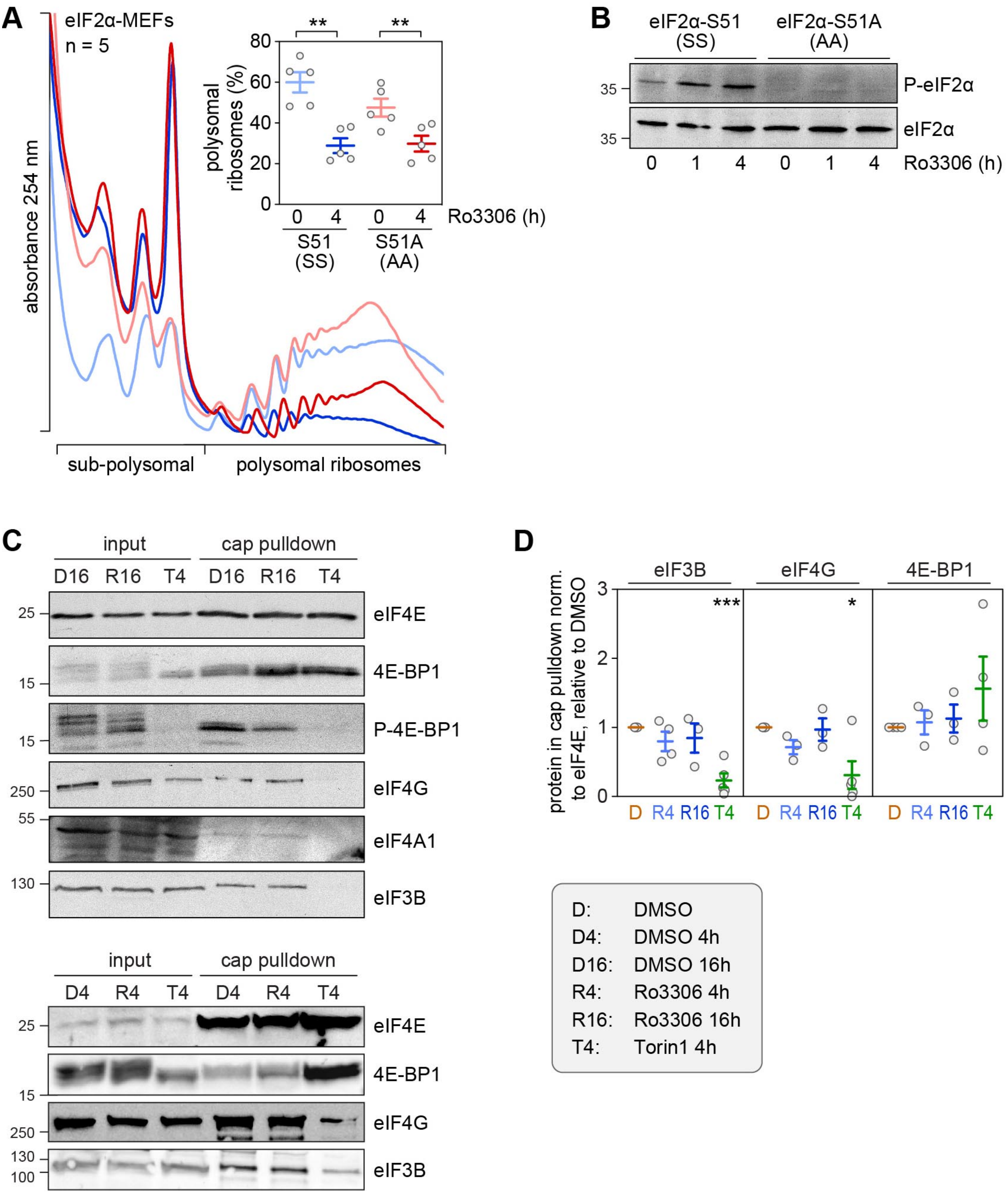


Figure S3. **Signaling pathways downstream of CDK1i. (A)** eIF2α WT S51 (SS) and phosphodeficient S51A (AA) MEFs were treated with DMSO or Ro3306 (10 µM) for 4h. Polysome profiles were recorded after sucrose density gradient centrifugation; the percentage of polysomal ribosomes is represented in the inset (average ± SEM, n = 5). Statistical significance was determined by paired Student's t test; **, $p \leq 0.01$. **(B)** The phosphorylation level of eIF2α (S51) was assessed in cytoplasmic lysates from samples in

(A) by Western blot analysis. **(C)** Cytoplasmic lysates from DMSO-, Ro3306- and Torin1-treated HeLa cells were subjected to cap pulldown experiments using 7-methyl-GTP agarose beads. Cap-associated factors eIF4E, 4E-BP1, P-4E-BP1 (T37/T46), eIF4G, eIF4A1 and eIF3B were detected by Western blot analysis. **(D)** For quantification of cap-associated factors, levels of eIF3B, eIF4G and 4E-BP in the cap pulldown were normalized to the level of precipitated eIF4E, and depicted relative to the value obtained for the DMSO control (average ± SEM, $n \geq 3$). Statistical significance was determined by one sample Student's t test; *$p \leq 0.05$, *** $p \leq 0.001$.

*Figure S4, Haneke et al.*

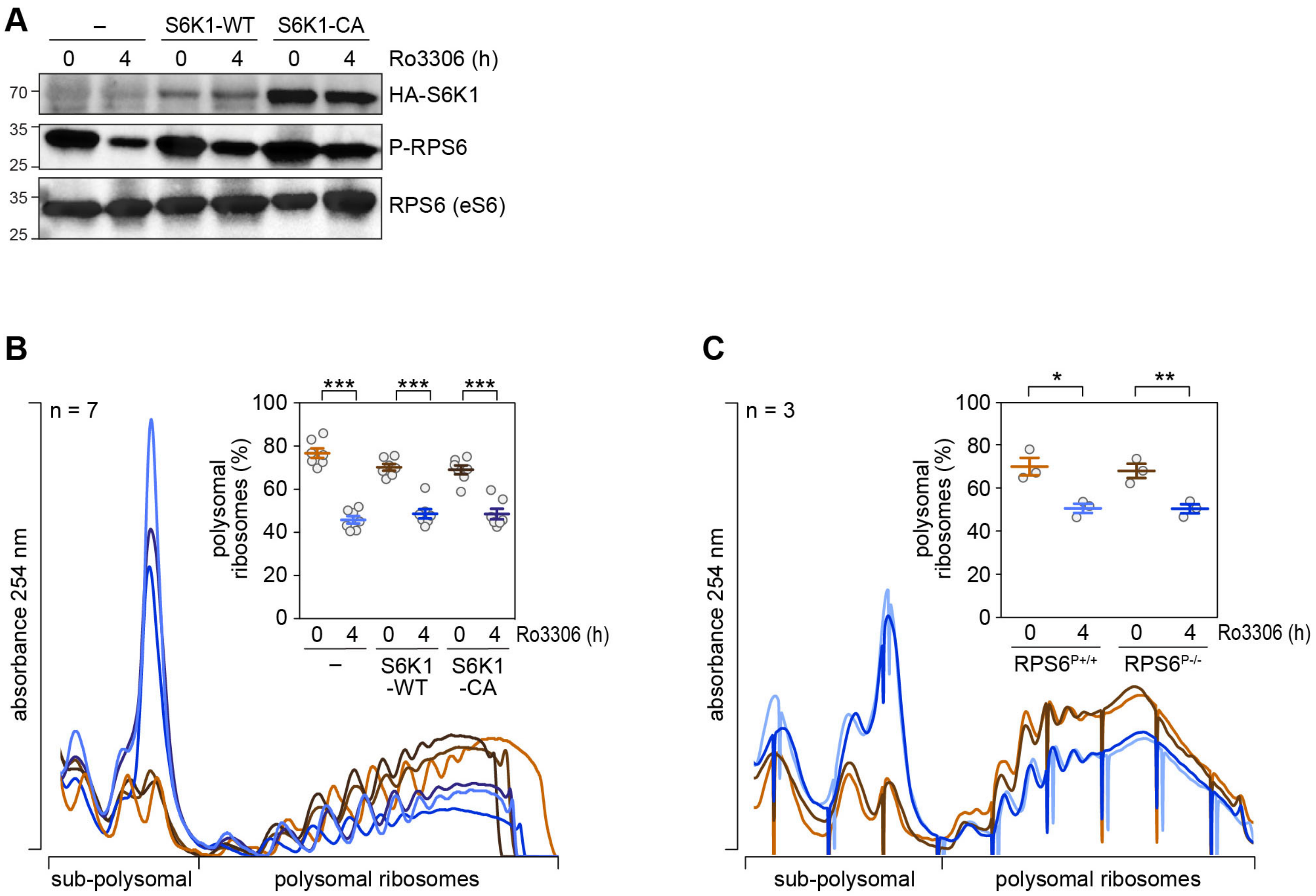


Figure S4. **Role of S6K1 and RPS6 phosphorylation in CDK1i-induced translation inhibition. (A)** HeLa cells were stably transduced with HA-S6K1-WT or constitutively active HA-S6K1-CA. Cells were treated with DMSO or Ro3306 (10 μM) for 4h. The levels of S6K1 overexpression and the phosphorylation status of RPS6 were assessed by Western blot analysis. **(B)** Polysome profiles were recorded from HeLa, HeLa-HA-S6K1-WT and HeLa-HA-S6K1-CA cells treated with either DMSO or Ro3306 (10 μM) for 4h; the percentage of polysomal ribosomes is represented in the inset (average ± SEM, n = 7). **(C)** Polysome profiles were recorded from RPS6 WT ($RPS6^{P+/+}$) and RPS6 phosphodeficient S235A, S236A, S240A, S244A, S247A ($RPS6^{P-/-}$) MEFs treated with either DMSO or Ro3306 (10 μM) for 4h; the percentage of polysomal ribosomes is represented in the inset (average ± SEM, n = 3). In (B and C), statistical significance was determined by paired Student's t test; *, $p \leq 0.05$; **, $p \leq 0.01$; ***, $p \leq 0.001$.

*Figure S5, Haneke et al.*

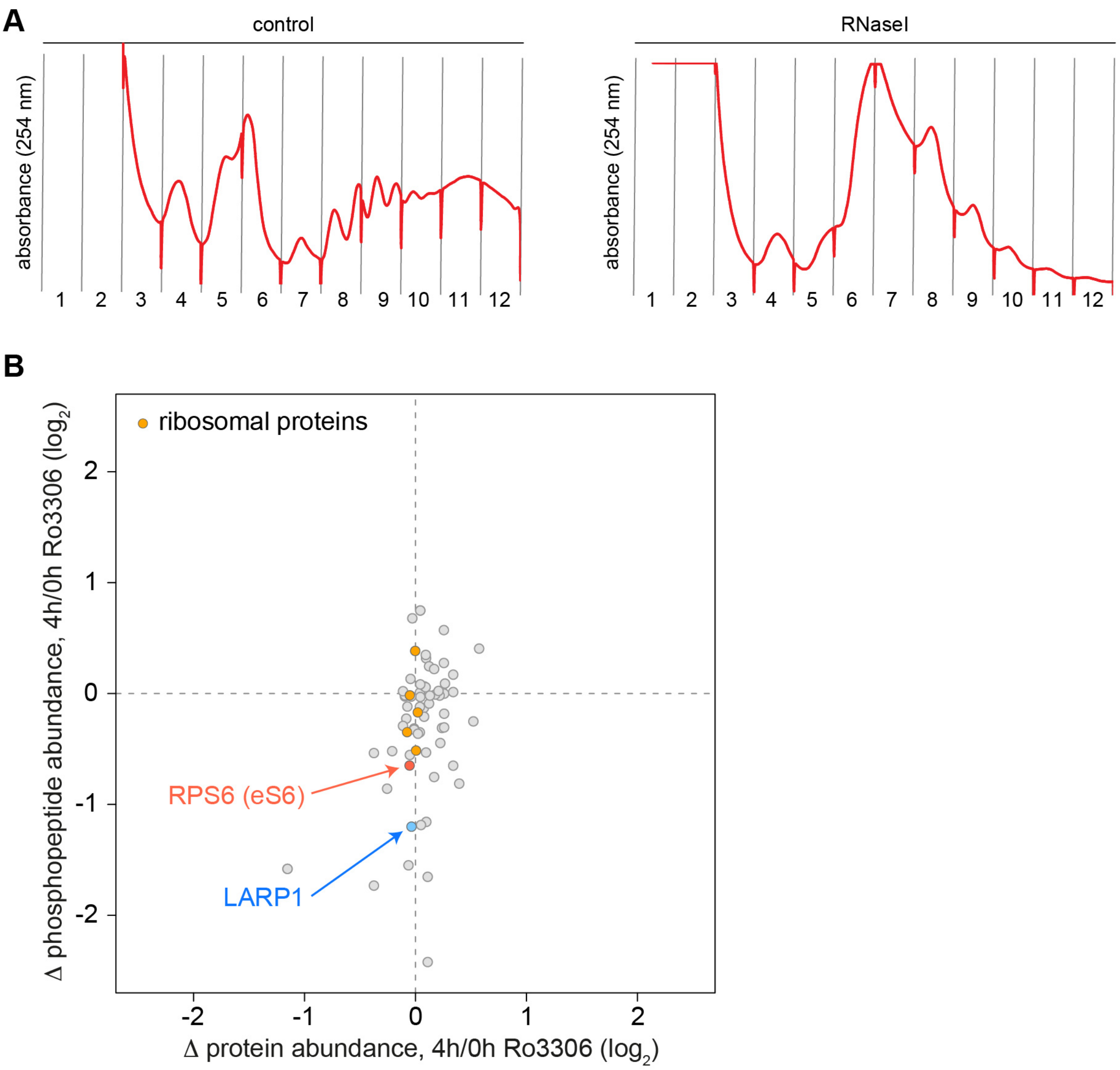


Figure S5. **Polysome control and CDK1-dependent phosphorylation events. (A)** Prior to sucrose density gradient centrifugation, HeLa cell lysates were subjected to RNase I digestion or left untreated for control. Absorbance at 254 nm was monitored during fractionation. **(B)** For phosphoproteomics of ribosomal fractions, HeLa cells were SILAC-labeled and either treated with DMSO (heavy) or Ro3306 (light) for 4 h. After lysis and disassembly of polysomes in low magnesium buffer, samples were mixed, and ribosomal fractions obtained by sucrose density centrifugation were subjected to Wessel-Flügge precipitation. Phosphopeptides were enriched by TiO2-SIMAC-HILIC (TiSH), fractionated and analyzed by mass spectrometry followed by MaxQuant analysis. For all phosphopeptides detected under both conditions, the ratio (Δ phosphopeptide abundance, 4h/0h Ro3306) was plotted against the ratio of the corresponding total protein (Δ protein abundance, 4h/0h Ro3306). Phosphopeptides derived from LARP1 (blue), RPS6 (red) and other translation regulators (orange) are color-coded.

*Figure S6, Haneke et al.*

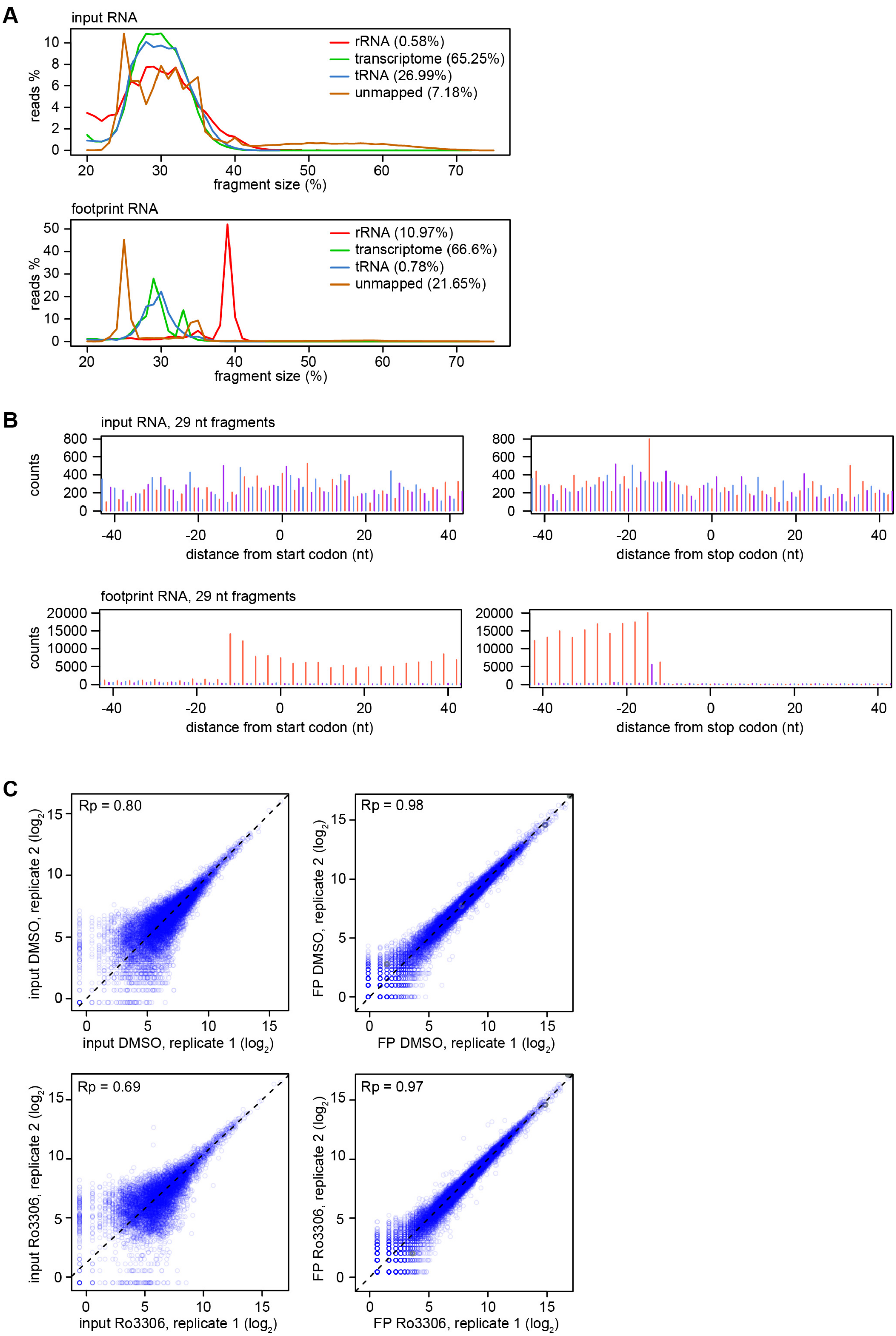

Figure S6. **Ribo-Seq quality assessment. (A)** All reads from the input RNA and ribosome footprint samples were mapped to human tRNA sequences, rRNA sequences, the human transcriptome (wgEncodeGencodeBasicV27) and a yeast transcriptome (sacCer3ensGene). Depicted is the fragment size distribution of the four groups (rRNA, tRNA, transcriptome, unmapped) from one representative input RNA (top) and footprint sample (bottom). **(B)** Reads from one representative input RNA (top) and footprint sample (bottom) were aligned at the 5' end, and the distribution of all 29 nt long fragments is depicted according to their position relative to the start codon (left panels) and to the stop codon (right panels). **(C)** To assess reproducibility between repeat experiments, two repeats are plotted against each other representative for each condition (input DMSO, input Ro3306, footprint DMSO, footprint Ro3306).

*Figure S7, Haneke et al.*

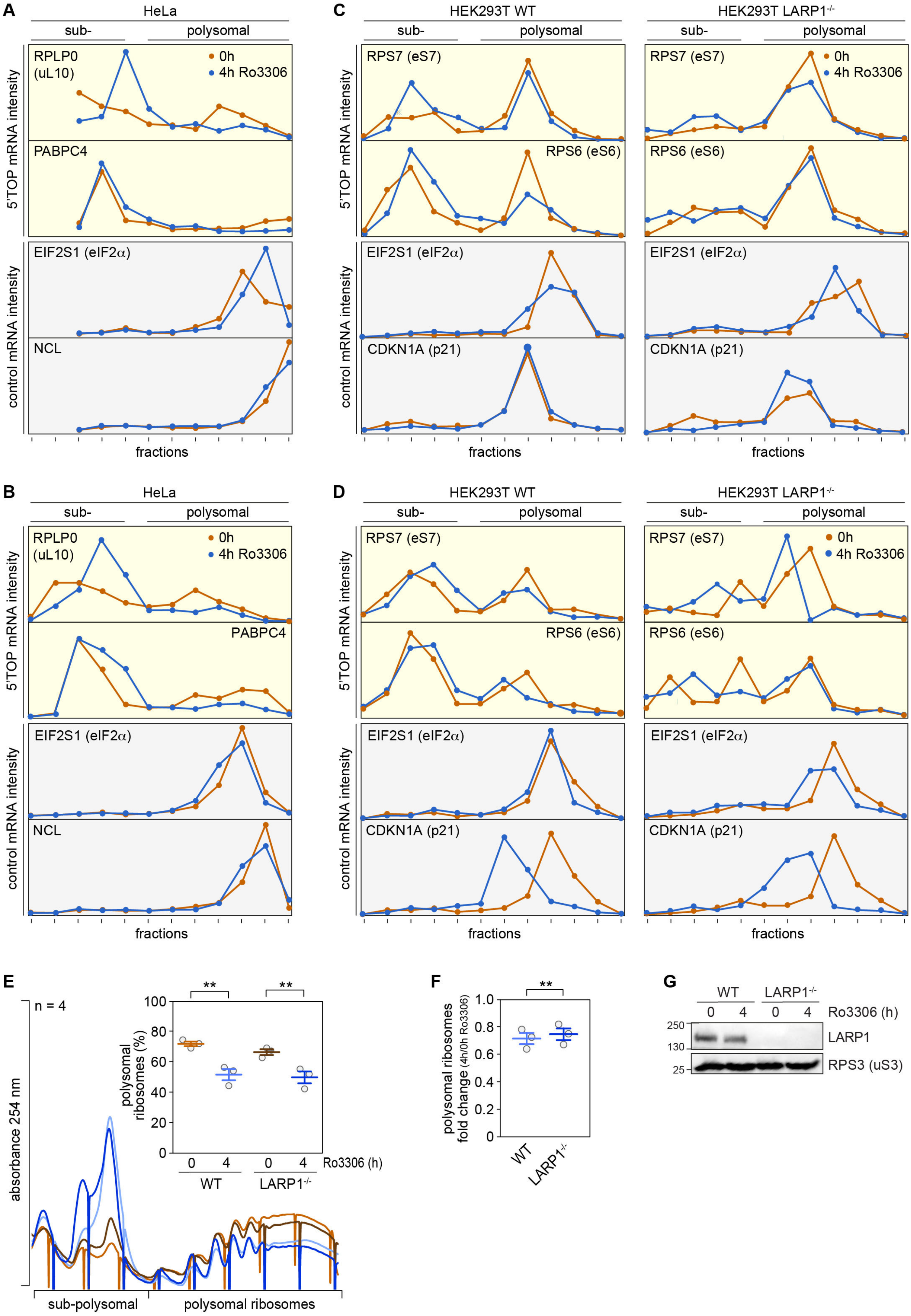

Figure S7. **Signaling pathways downstream of CDK1i. (A, B)** Polysome association of 5'TOP (RPLP0 and PABPC4) and ORF size-matched non-TOP (EIF2S1 and NCL) mRNAs was analyzed by polysome fractionation and subsequent qPCR analysis from DMSO- or Ro3306-treated (10 μM, 4 h) HeLa cells. **(C, D)** Polysome association of 5'TOP (RPS6 and RPS7) and ORF size-matched non-TOP (EIF2S1, CDKN1A) mRNAs was analyzed by polysome fractionation and subsequent qPCR analysis from DMSO- or Ro3306-treated (10 μM, 4 h) HEK293T WT or LARP1$^{-/-}$ cells. **(E)** Polysome profiles were recorded from HEK293T WT or LARP1$^{-/-}$ cells treated with either DMSO or Ro3306 (10 μM) for 4h; the percentage of polysomal ribosomes is represented in the inset (average ± SEM, n = 3). **(F)** The fold-change in polysomal ribosomes (4 h Ro3306 / DMSO control) was calculated based on polysome profiles in (E). In (E-F), statistical significance was determined by paired Student's t test; **, $p \leq 0.01$. **(G)** LARP1 expression was monitored in HEK293T WT and LARP1$^{-/-}$ cells by Western blot analysis, RPS3 serves as loading control.
